# The Enterococcal Polysaccharide Antigen: from structure to biosynthesis and function

**DOI:** 10.1101/2024.06.26.600781

**Authors:** Jessica L Davis, Joshua S Norwood, Robert E Smith, Finn O’Dea, Krishna Chellappa, Michelle L Rowe, Mike P Williamson, Graham P Stafford, Evguenii Vinogradov, Emmanuel Maes, Yann Guérardel, Stéphane Mesnage

**Affiliations:** School of Biosciences, University of Sheffield, Sheffield, UK; School of Clinical Dentistry, University of Sheffield, Sheffield, UK; Vaccine and Emerging Infections Research, Human Health Therapeutics Research Centre, National Research Council, Ottawa, Canada; University of Lille, CNRS, Inserm, CHU Lille, Institut Pasteur de Lille, US 41 - UAR 2014 - PLBS, Lille, France; UMR 8576 - UGSF - Unité de Glycobiologie Structurale et Fonctionnelle, CNRS, Université de Lille, Lille, France; Institute for GlycO-Core Research (iGCORE), Gifu University, Gifu, Japan

**Keywords:** Enterococcal Polysaccharide Antigen, L-Rhamnose, polysaccharide structure, Bacteriophages, innate immune evasion

## Abstract

L-Rhamnose-containing polysaccharides are produced by Streptococci and Enterococci. They define Lancefield serotypes and represent promising candidates for the design of glycoconjugate vaccines. The Enterococcal Polysaccharide Antigen produced by the opportunistic pathogen *Enterococcus faecalis* plays a critical role in normal growth, division, biofilm formation, antimicrobial resistance, phage susceptibility, and innate immune evasion. Despite the critical role of this polymer for *E. faecalis* physiology and host-pathogen interactions, little information is available on its structure and biosynthesis. Here, we elucidate the structure of the intact EPA produced by *E. faecalis* OG1RF. We report the structure of the linkage unit, revealing an unprecedented complexity of the rhamnose backbone and decorations. Finally, we explore the impact of several EPA structural modifications on innate immune evasion and recognition by bacteriophages. This work represents a first step towards the functional characterisation of EPA for the rational design of therapeutic strategies against a group of important pathogens.

## Main

Cell wall polysaccharides containing L-rhamnose are commonly displayed on the cell surface of Gram-positive ovococci and their structural diversity underpins the definition of Lancefield serotypes^1^. L-rhamnose polysaccharides are essential for normal cell growth, division, antimicrobial resistance, and pathogenesis^2,3^ and have been proposed as targets for the development of glycoconjugate vaccines^4,5^. The work on rhamnose-containing carbohydrates has largely focused on the Group A carbohydrate (GAC), produced by *Streptococcus pyogenes* (group A Streptococcus, GAS)^6^ and the serotype C carbohydrate (SCC) produced by *Streptococcus mutants*^7,8^. The structural analysis of the GAC elucidated the mechanisms that confer resistance to human innate immune effectors such as group II phospholipase A2 and cationic antimicrobial proteins^9,10^. Several studies have also shed light on the structure of polysaccharides containing L-rhamnose in Lactococci. In this group of bacteria, genomic and structural studies have defined four major groups of polymers (A-D) and established their contribution to cell surface recognition by bacteriophage^11-13^

Most Enterococci, including the human pathogens *Enterococcus faecalis* and Enterococcus faecium, produce a complex surface polymer containing L-rhamnose called the Enterococcal Polysaccharide Antigen^1,14,15^ defining the Lancefield serotype D. In *E. faecalis*, the production of EPA is required for normal cell growth, and biosynthetic mutants display morphological defects^16,17^. EPA has also been identified as a key polymer for biofilm formation^17-19^, antimicrobial resistance^20,21^, cell surface recognition by bacteriophage^16,22^ and virulence^23^. The presence of EPA at the cell surface of *E. faecalis* promotes host colonization^2421^ and prevents phagocytosis^14,25-27^.

The structural and functional characterization of EPA is extremely challenging. This polymer is encoded by two adjacent chromosomal loci (fig. 1a). One locus consists of genes *epaA* to *epaQ*, conserved across *E. faecalis* and encoding the synthesis machinery of the EPA “rhamnose backbone”. The other locus, downstream of *epaQ* contains *epaR* and a variable number of genes (between 10 and 20) depending on the strain considered. This second region encodes so-called EPA “decorations”. Recent work clearly established that EPA decorations are essential for the biological activity of this polymer^27^. The complete structure of EPA, and the structural properties required for innate immune evasion and other processes, however, remain unknown.

**Fig. 1.**
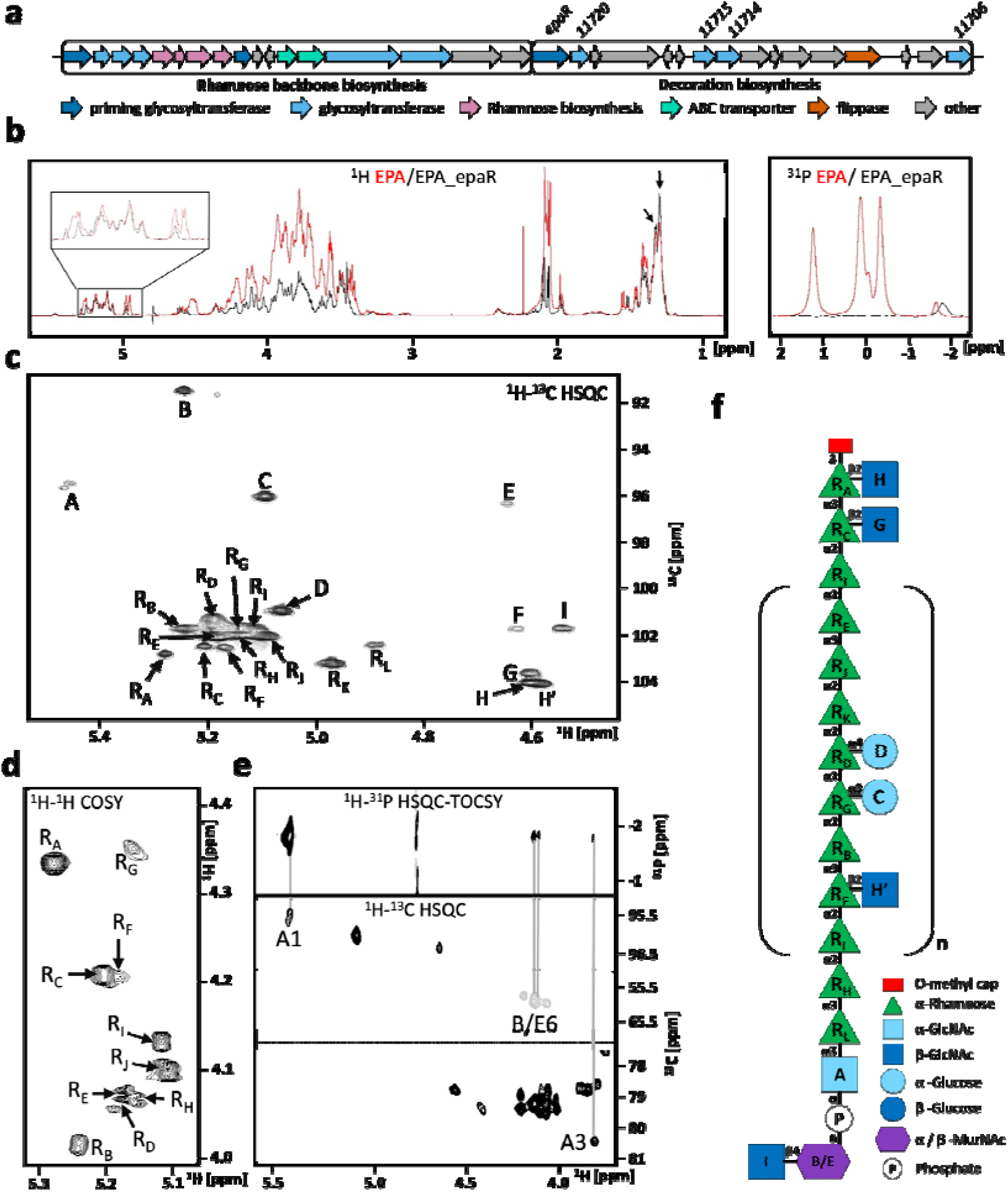
Structure of the EPA rhamnose backbone (EPA_epaR product). **a**, The EPA biosynthetic locus. **b**, 1D ^1^H (left) and ^31^ P NMR analysis of EPA purified from *E. faecalis* OG1RF (red), and ΔepaR mutant (black, EPA_epaR). Analysis of α-anomeric (boxed) and rhamnose methyl (arrows) signals suggests that EPA_epaR contains fewer monosaccharide spin systems as compared to EPA, but an equal amount of rhamnosyl residues. **c**, Anomeric region of the EPA_epaR^1^ H-^13^C HSQC spectrum. 22 monosaccharides were detectable (A-I, H’, and R_A_-R_L_). **d**, R_A_-R_J_ H1/H2 region of the EPA_epaR ^1^ H-^1^H COSY spectrum. The unique H1/H2 shifts of R_A_-R_J_ enabled the separation of each spin system. e, ^1^H-^31^P HSQC-TOSCY spectrum (top panel) of EPA_epaR. Signals could be mapped (vertical lines) onto the ^1^H-^13^C HSQC spectrum (bottom three panels), enabling the characterisation of the EPA_epaR-peptidoglycan linkage. **f**, Complete structure of the EPA rhamnose backbone (EPA_epaR).

The only structural information regarding EPA concerns fragments produced by acid treatment of the polymer from *E. faecalis* V583^28^. The rhamnose (Rha) backbone was described as a rhamnan chain made of a repeating unit of six residues linked via α-1,2 and α−1,3 bonds and substituted by α-glucose (Glc) and β-*N*-acetylglucosamine (GlcNAc) residues at positions C-2 and C-3, respectively. Two decoration subunits were identified, containing α/β-Glc, β-*N*-acetylgalactosamine (GalNAc), α-Rha and ribitol-phosphate groups, typically found in teichoic acids of Gram-positive bacteria^2^. A biosynthetic pathway was also proposed based on sequence analysis of the conserved and variable epa loci. However, due to the experimental approach used for the structural analysis, no information is available concerning the architecture of the intact polymer and most of the roles assigned to epa genes remained hypothetical.

In this study, we elucidate the structure of the intact EPA polymer produced by *E. faecalis* and reveal the unprecedented complexity of this macromolecule. Furthermore, we describe the structure of EPA produced by several decoration mutants and test the contribution of specific structural motifs in the evasion of phagocytosis and recognition by bacteriophages.

## Results

### EPA is a complex polymer with multiple phosphate-linked subunits

We previously reported the structure of EPA fragments produced by acid treatment of the full-length *E. faecalis* polymer in strain V583^28^. Although this represented an important step for the structure/function analysis of EPA, this work did not provide information about the entire polymer architecture or minor modifications, which may be essential for the biological activity of EPA. We sought to investigate the architecture of the intact EPA polymer in *E. faecalis* OG1RF, a clinical strain used for virulence studies^29^. NMR analysis of EPA revealed an unprecedented complexity, identifying over 25 unique monosaccharide spin systems (supplementary fig. 1a), as well as characteristic shifts corresponding to glycerol-phosphate groups (supplementary fig. 1b). 1D ^31^P NMR confirmed the presence of several phosphate groups within EPA, resonating at 1.2 ppm, 0.11 ppm, 0.07 ppm, -0.34 ppm, and -1.65 ppm (fig. 1b, right). Based on signal intensity and their respective chemical shifts, we hypothesised that EPA contains several distinct building blocks and possibly a phosphodiester linkage unit to peptidoglycan (most likely the lowest intensity peak at -1.65 ppm). The two peaks at 0.11 ppm and 0.07 ppm have very similar resonances but distinct relative abundances. This suggests that these two phosphate groups exist in very similar, but not identical, environments most likely corresponding to major/minor structural variants of EPA.

### The EPA backbone is an *O*-methyl capped undecasaccharide

To reduce the complexity of EPA, we used *epa* glycosyltransferase mutants that produce truncated versions of the full-length polymer^27^. We first decided to use an *epaR* mutant^22^. This gene is conserved across strains and predicted to encode a priming glycosyltransferase^16^ potentially responsible for the first committed step of decoration biosynthesis. As anticipated, EPA purified from the ΔepaR mutant did not contain any EPA decoration signals previously reported^27^ but retained the same proportion of rhamnose methyl signals (fig. 1b, left). 1D phosphorus NMR confirmed the absence of EPA decoration subunits, with only one weak phosphorus signal detected (fig. 1b, right) most likely corresponding to the EPA-peptidoglycan linkage group.

A total of 22 monosaccharide spin systems were identified by ^1^H-^13^C HSQC NMR (fig. 1c). The majority of EPA_epaR signals corresponded to rhamnose (Rha) residues (R_A_-R_L_). Alpha-glucose (Glc; residues C and D), α-*N*-Acetylglucosamine (GlcNAc; residue A) and β-GlcNAc (residues F, G, H, and H’) were also identified, as well as signals corresponding to the peptidoglycan linkage unit (residues B, E, and I; supplementary fig. 2). Rhamnose shifts R_b_-R_j_ were virtually impossible to distinguish due to overlapping anomeric carbon/proton signals in the ^1^H-^13^C HSQC. To deal with this issue, spin systems were differentiated using the shifts of protons one and two, recorded using a ^1^H-^1^H COSY experiment (fig. 1d). Using the anomeric shifts as reference, all proton and carbon shifts for each monosaccharide spin system were assigned (supplementary Table 1) and all glycosidic linkages were resolved (supplementary Table 2) using a series of 2D NMR experiments (^1^H-^13^C HSQC, ^1^H-^1^H COSY, ^1^H-^1^H TOSCY, ^1^H-^13^C HSQC-TOCSY, ^1^H-^13^C HMBC and ^1^H-^1^H NOESY). No NOE signals were detected for residue F, most likely caused by the low prevalence of this spin system.

**Fig. 2.**
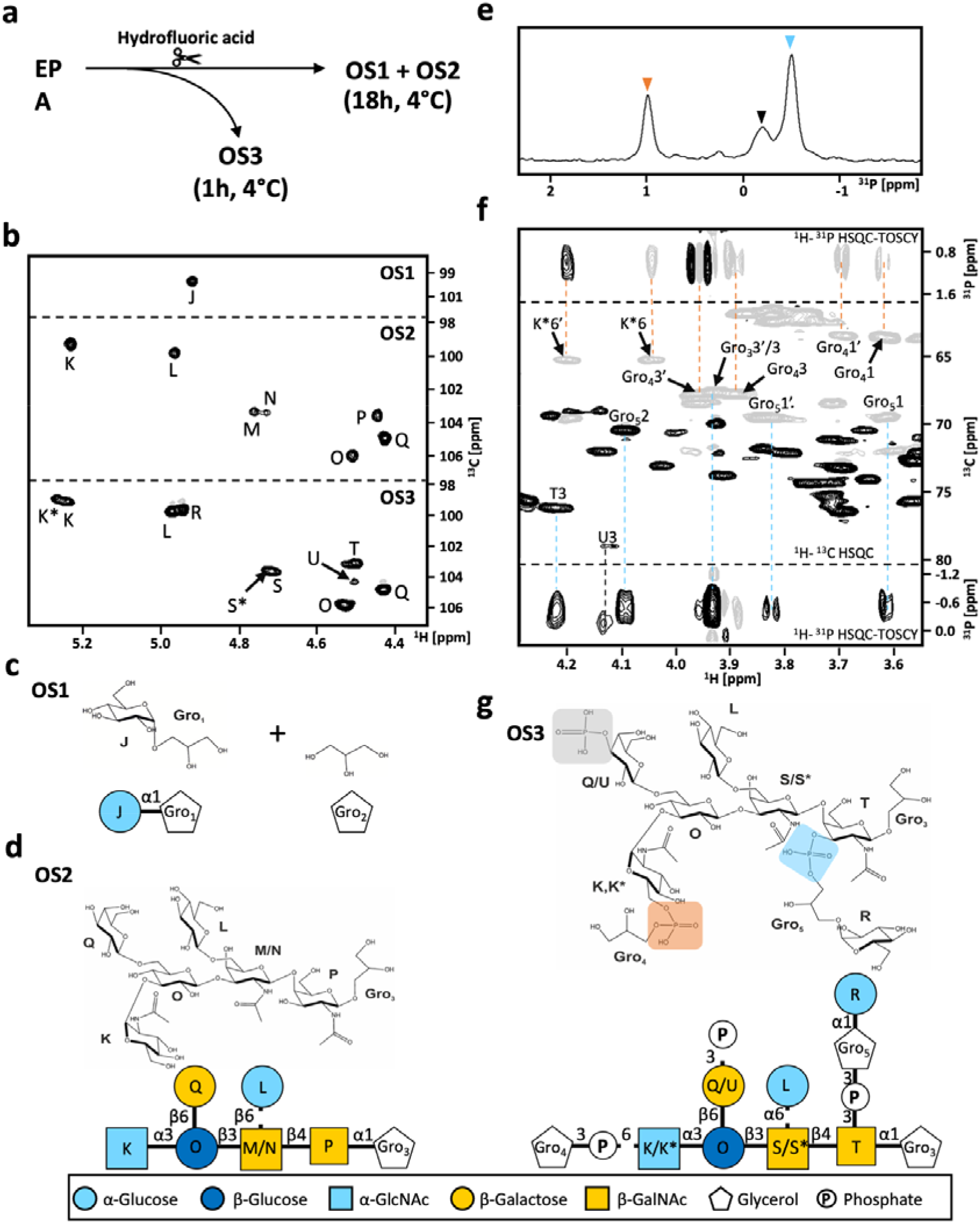
EPA decoration structure. **a**, Schematic of EPA treatment with hydrofluoric (HF) acid to produce OS1-OS3. **b**, Anomeric regions of the ^1^H-^13^C HSQC spectra acquired on OS1-OS3. Structure of fragments **c**, OS1 and **d**, OS2. **e**, OS3 ^31^P NMR spectrum. **f**, Assignment of phosphodiester linkage units in OS3. **g**, OS3 structure.

To complete the structure of EPA_epaR, a ^1^H-^31^P HSQC-TOCSY spectrum was recorded (fig. 1e). It revealed that EPA is linked to peptidoglycan by a phosphodiester group bound to the anomeric position of α-GlcNAc (residue A) and the sixth position of MurNAc (residues B/E; supplementary fig. 2).

Across this study, all EPA samples were purified enzymatically with mutanolysin, leaving a monomeric peptidoglycan scar which contains a MurNAc residue existing in both an α and β form. To identify the anomeric ^1^H-^13^C-HSQC signals produced by peptidoglycan fragments, cell wall sacculi were purified from *E. faecalis* OG1RF and digested with mutanolysin. Monomeric peptidoglycan fragments were purified by size exclusion chromatography and a ^1^H-^13^C-HSQC 2D NMR experiment was recorded (blue). The anomeric region was overlayed with the ^1^H-^13^C-HSQC spectrum of EPA_epaR sample. Residues B, E and I (bold) have identical shifts in both samples and must therefore correspond to the three anomeric signals (α-MurNAc, β-MurNAc and β-GlcNAc) from the peptidoglycan scar.

The structure of the EPA rhamnose backbone, described in fig. 1f, highlights the intricacies and original properties of this polymer. It consists of three distinct sections: a peptidoglycan linkage unit, (ii) a polymerized unit, and (iii) a capping unit. The linkage unit is composed of a phosphodiester bond connecting the α-GlcNAc (residue A) to the sixth position of β-MurNAc within the peptidoglycan layer. Two α-Rha residues (residues R_H_ and R_L_) substitute this α-GlcNAc to make the acceptor structure for the polymerisation of the repeating unit. The polymeric unit is composed of an octameric linear Rha chain linked via α-1,2 and α-1,3 glycosidic bonds (residues R_I_, R_F_, R_B_, R_G_, R_D_, R_K_, R_J_ and R_E_). This repeat is consistently substituted with one β-GlcNAc residue and two α-Glc residues at positions 2, 3 and 4 corresponding to Rha residues R_F_, R_G_ and R_D_, respectively. The polymerisation of this unit is terminated by the addition of Rha residues R_c_ and R_A_ onto residue R_I_. Residue R_A_ is unique because it is substituted at the second and third positions with β-GlcNAc and an *O*-methyl group respectively, preventing further polymerisation through α-1,2 and α-1,3 bonds. This final residue therefore acts as a capping unit that terminates the polymerisation of the Rha chain and potentially represents a structural cue for transport and/or the addition of the decoration subunits. By comparison of relative signal intensities within the ^1^H-^13^C HSQC, corresponding to residues within and outside of the octameric repeat, we estimate that this *O*-methyl group is added after approximately 3-4 repeats of the octameric rhamnose unit.

### EPA decorations are branched and contain heterogeneous modifications

Phosphate NMR analysis of EPA suggests that only decoration subunits contain phosphodiester bonds (fig. 1b). This feature allowed us to generate decoration fragments from EPA after treatment with hydrofluoric acid (fig. 2a). Incubation times of 1 and 18 hours released three fragments with distinct complexity (OS1-OS3), as confirmed by ^1^H-^13^C HSQC (fig. 2b). Treatment for 18 hours was predicted to cleave all phosphodiester bonds, and as expected, only one phosphorus signal (corresponding to free phosphate) was detected across OS1 and OS2 (supplementary fig. 3). The structures of OS1 and OS2 were solved using the same repertoire of NMR experiments previously described. All assigned shifts and information regarding residue connectivity are described in supplementary Tables 3 and 4.

**Fig. 3.**
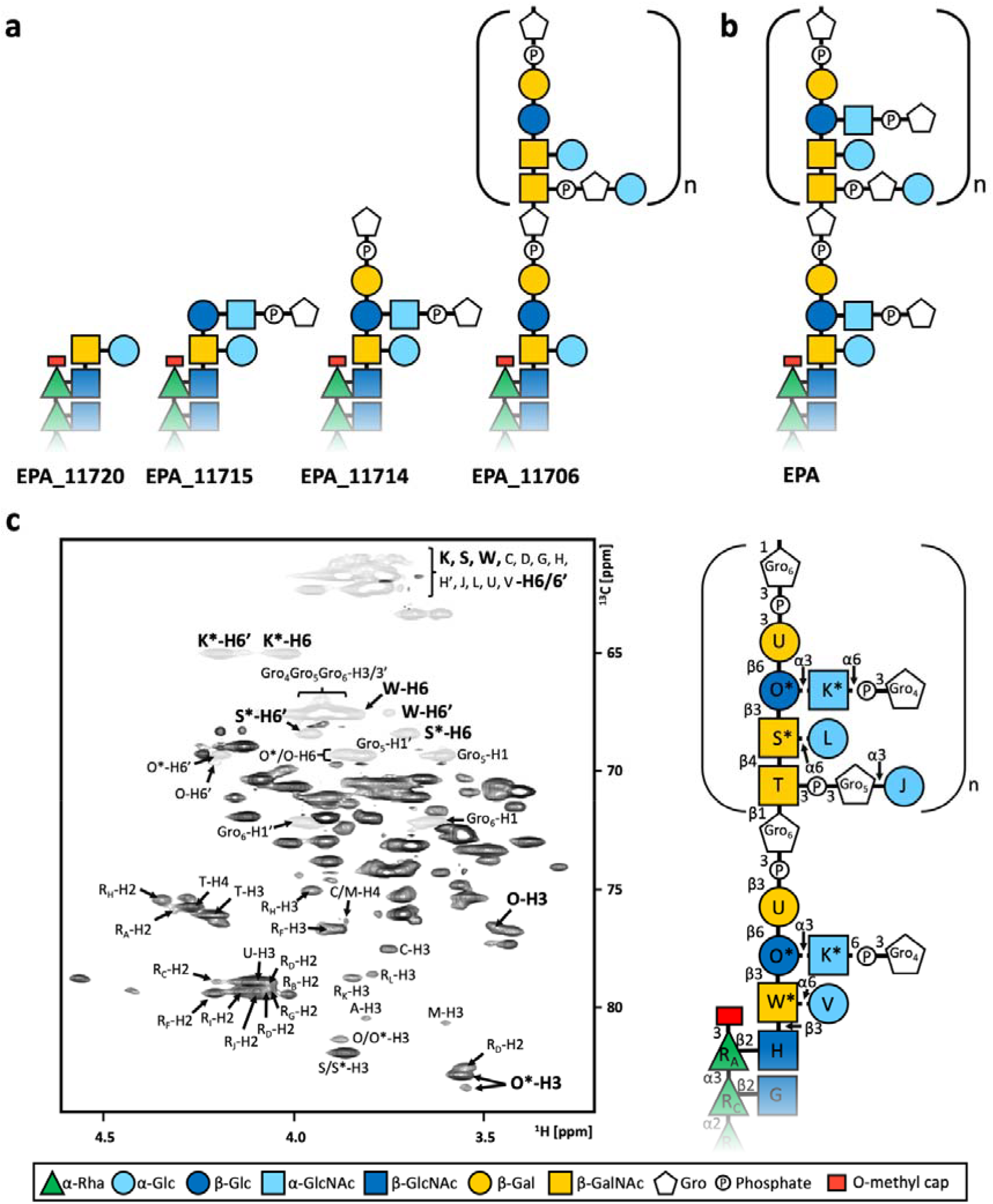
The complete structure of EPA. **a**, Structure of EPA purified from mutants Δ*11720*, Δ*11715*, Δ*11714* and Δ*11706* (from left to right). **b**, The structure of EPA. In all cases **(a, b)** the EPA rhamnose backbone was found to be identical to EPA_epaR (fig. 1f). **c**, (left) ring system region of the EPA ^1^H-^13^C HSQC spectrum with shifts corresponding to atoms involved in EPA residue connectivity. Some residues exist in two forms (bold); substituted (marked with an asterisk) or unsubstituted. This is reflected in the final structure of EPA, specifically for residues K, O and W/S which are inconsistently substituted at the sixth, third and sixth positions, respectively.

OS1 contained two separate species: free glycerol and an α-Glc(1-3)Glycerol (fig. 2c). In contrast, OS2 was composed of a single more complex, branched, oligosaccharide, made of α-Glc (residue L), α-GlcNAc (residue K), β-Glc (residue O), β-Gal (residue Q), β-GalNAc (residues M/N and P) and glycerol (fig. 2d). One β-GalNAc residue was found in two forms, unsubstituted (residue N) or substituted (residue M) by an α-Glc (residue L) at the sixth position. This suggests that EPA decorations are heterogeneous and can be modified with terminal α-Glc residues.

OS3 was purified following partial acid cleavage (1-hour treatment) and revealed the connection between OS1 and OS2 in the intact decoration structure. The full assignment of OS3, and connectivity between monosaccharides and aglycon groups (i.e. phosphate and Gro), were determined by NMR analysis and are described in supplementary Tables 5 and 6. OS3 retains three phosphodiester bonds with signals at 0.98 ppm, -0.21 ppm, and -0.51 ppm (fig. 2e), linking the OS1 fragments to OS2, as resolved by ^1^H-^31^P HSQC-TOCSY (fig. 2f). From the phosphorus shift at 0.98 ppm, magnetisation transfer was observed to the sixth position of α-GlcNAc (residue K* in OS3, residue K in OS2) and the proton 1 and 3 shifts of glycerol (residue Gro_4_). The carbon 1 and 3 shifts of glycerol Gro_4_ are at 63 and 67 ppm, respectively, and it was therefore concluded that this glycerol corresponds to the free glycerol (residue Gro_2_) isolated in OS1. The phosphorus shift at -0.51 ppm exhibited magnetisation transfer to the third position of β-GalNAc (residue T in OS3, residue P in OS2) and to the proton 1, 2 and 3 shifts of glycerol (residue Gro_5_). As the carbon 1 and 3 shifts of glycerol Gro_5_, are at 69 and 67 ppm, respectively, it was concluded this glycerol corresponds to glycerol Gro_1_ within OS1. The minor phosphorus shift at -0.21 ppm only showed magnetisation transfer to the third position of β-Gal (residue U in OS3, residue Q in OS2) and as such is most likely a phosphomonoester.

The final structure of OS3 is shown in fig. 2g. Collectively, NMR analysis indicated that EPA decorations have a branched structure, made of 3 subunits linked by phosphodiester bonds. The structure of the intact monomeric decoration unit suggested that EPA decorations are most likely polymerized through the β-galactose (residue H). Interestingly the heterogenous substitution of one of the β-GalNAc residues (residues M/N in OS2 and S/S* in OS3) with a terminal α-Glc was present in both OS2 and OS3 and indicate the presence of minor modifications within EPA decorations.

### EPA decorations are anchored onto the rhamnose backbone capping unit

To explore how the rhamnose backbone and EPA decorations are assembled to form the final polymer, we elucidated the structure of EPA produced by mutants in glycosyltransferase genes encoded within the decoration locus (fig. 1a). Three mutants (Δ*11720*, Δ*11715*, Δ*11714*) were already available^27^ whilst the in-frame deletion in gene *11706* was built here. We sequentially solved the structure of EPA produced by these four mutants (fig. 3a). Based on the increasing complexity of anomeric regions in ^1^H-^13^C HSQC spectra, we started with EPA_11720, followed by EPA_11715, EPA_11714 and EPA_11706 (supplementary fig. 4). In each case, 1D ^1^H, ^31^P, as well as 2D ^1^H-^1^H, ^1^H-^13^C and ^1^H-^31^P-NMR experiments were recorded and used to assign new residues and sequentially resolve the structure of each polysaccharide (supplementary Tables 7-14).

**Fig. 4.**
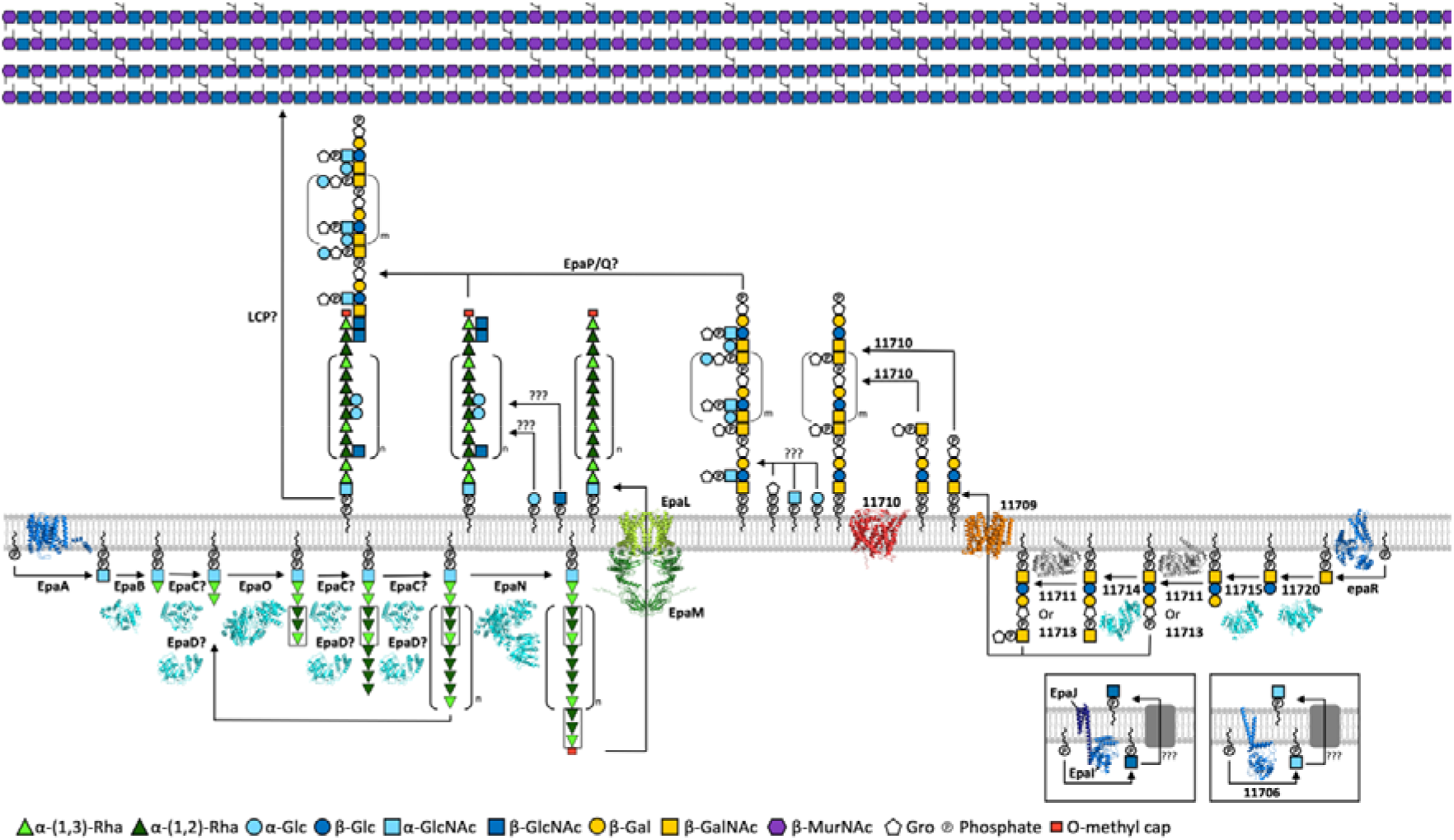
Model of EPA biosynthesis. The AlphaFold-2^35^ and AlphaFold-Multimer ^36^ structures of the priming glycosyltransferases (EpaA/EpaR/EpaI:J/11706; dark blue), glycosyltransferases (EpaB-D/EpaN-O/11720/11715-14; light blue), glycerol-phosphate transferases (11713/11711; grey), the ABC transporter complex (EpaL/EpaM; green), the flippase (11709; orange), and the *O*-antigen ligase like protein (11710; red) thought to be involved in EPA biosynthesis are shown. Question marks denote biosynthetic steps where no enzyme has yet been assigned. Areas of symmetry within the EPA rhamnose backbone are highlighted (black boxes).

In all four mutants, the structure of the EPA rhamnose backbone remained the same (supplementary fig. 5). EPA_11720 (fig. 3a) revealed that EPA decorations are linked to the rhamnose backbone via the terminal β-GlcNAc (residue H) of EPA_epaR (fig. 1f). EPA_11720 is almost identical in structure to EPA_epaR but contains two additional monosaccharides: a β-GalNAc that substitutes residue H at position 3, and an α-Glc that substitutes β-GalNAc at position 6. In EPA_11715, the β-GalNAc is further substituted at position 3 by β-Glc, extending EPA decoration. This β-Glc is also substituted at position 3 by an α-GlcNAc, phosphorylated at position 6 by a phospho-glycerol group. Comparing EPA_11714 to EPA_11715 indicated that a β-Gal residue is connected to the β-Glc of EPA_11715, via a 1-6 bond and substituted by a phospho-glycerol group at position 3.

**Fig. 5.**
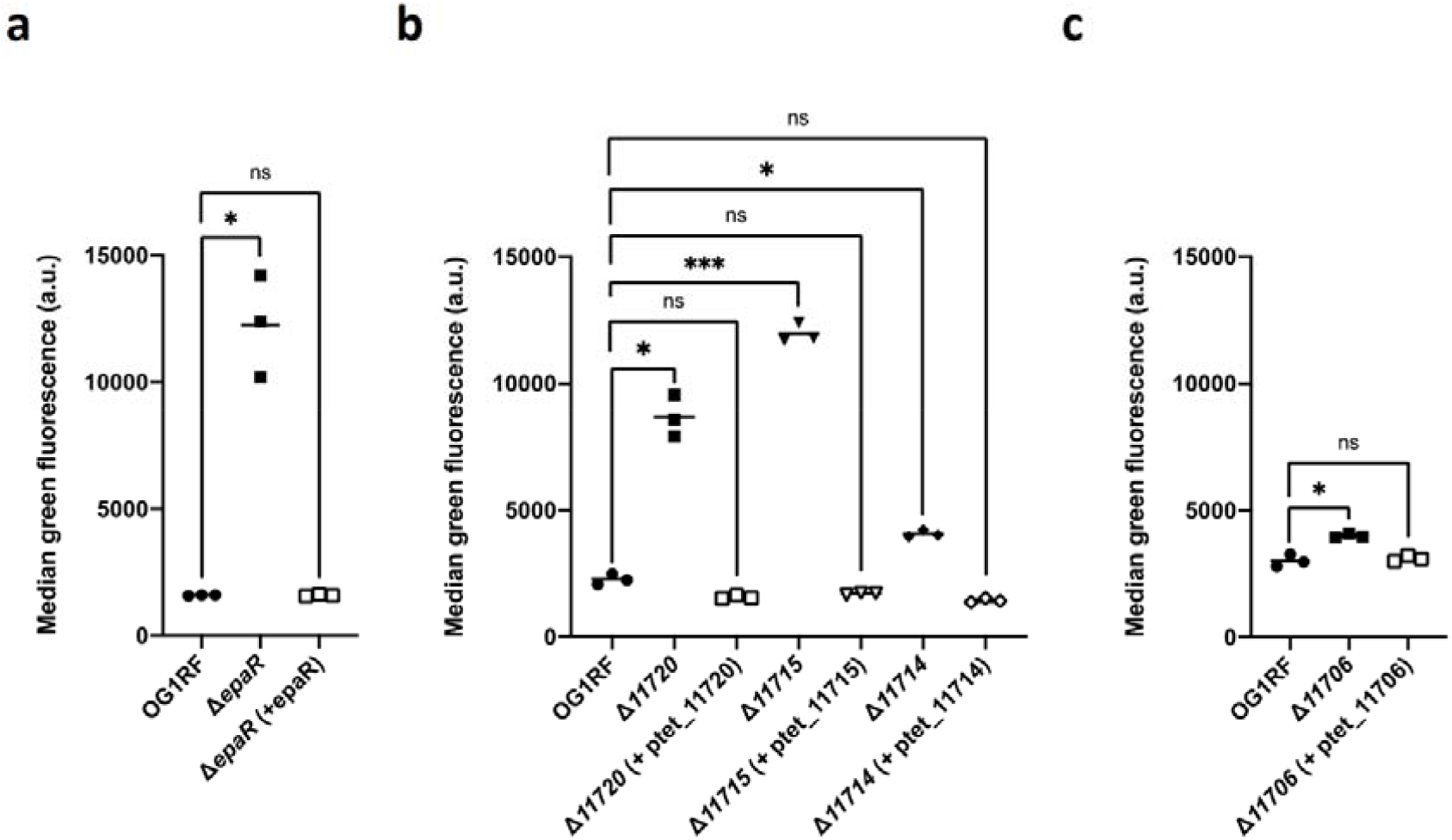
Contribution of EPA decorations to innate immune evasion. The ability of *E. faecalis* OG1RF isogenic deletion mutants (closed squares, triangles, and rhombus) to avoid uptake by iBMDMs as compared to WT cells (closed circles) was assessed using a phagocytosis assay. **a**, ΔepaR versus OG1RF (*P*=0.0284; *), **b** Δ*11720* versus OG1RF (*P*=0.0399; *), Δ*11715* versus OG1RF (*P*=0.004;***), Δ*11714* versus OG1RF (*P*=0.0108;*). **c**, Δ*11706* versus OG1RF (0.0307;*). All mutants exhibited increased uptake by iBMDMs and in all cases, complementation (open symbols) successfully restored the WT phenotype. NS, not significant; a.u., arbitrary units.

The structure of EPA_11706 confirmed that EPA decorations are polymerised through a phospho-glycerol group, built upon the Glycerol-(3-P-3)-β-Gal of EPA_11714. In EPA_11706, the glycerol group Gro_6_, is substituted at the first position by β-GalNAc, corresponding to the beginning of the next decoration subunit. This repeating structure is identical to OS3 (fig. 2g), except for the absence of the Glycerol-(1-P-6)-α-GlcNAc-(1-3) side chain substituting OS3 residue O, indicating that 11706 is essential for this modification.

Combining the information from EPA_11714 and EPA_11706 enabled us to solve the structure of the entire EPA polymer (fig. 3b). Based on the integration of signal intensities, the repeating EPA decoration unit appears to contain approximately 8-9 repeating units (OS3; fig. 2g) per EPA molecule. EPA residues were therefore named in accordance with OS3. Whilst the structure of the rhamnose backbone showed no heterogeneity in all mutants and WT EPA, the decoration structures consistently showed the same minor variations as observed in OS3 (fig. 2g) where β-glucose (residue O/O*), α-GlcNAc (residue K/K*) and β-GalNAc (residues W/W* and S/S*) exist in two forms (fig. 3c).

### EPA is assembled from two polymers produced independently

Based on the structure of EPA and bioinformatic analysis, we established a biosynthetic pathway defining the contribution of individual epa genes to EPA biosynthesis (fig. 4). Due to the presence of two separate transport systems within the EPA biosynthetic locus (fig. 1a), we propose that the rhamnose backbone and decorations are produced separately before being combined into a single polymer on the outer leaflet of the membrane and anchored to peptidoglycan.

Bioinformatic analysis of *epaA-epaQ* (supplementary Table 15), which encode the enzymes responsible for the production of the rhamnose backbone (fig. 1a), identified two priming glycosyltransferases (EpaA and EpaI/J), five rhamnosyltransferase candidates (EpaB, EpaC, EpaD, EpaN and EpaO), an ABC transporter complex (EpaL and EpaM)^16,30^, and two potential oligosaccharide transferases with GT-C type folds (EpaP and EpaQ). The presence of an ABC transporter suggests that the rhamnose chain is polymerised inside the cytoplasm^31,32^, whilst the α-Glc and β-GlcNAc substitutions (residues C, D, G, H and H’, fig. 1f) are likely to be added after translocation to the outside of the cell, analogous to the S. pyogenes rhamnopolysaccharide^33^. EpaI exhibits GlcNAc-phosphate-undecaprenol synthase activity which is enhanced by EpaJ. Here, our bioinformatic analysis proposes that EpaJ most likely acts as a recruiter protein, anchoring EpaI to the cell membrane, enabling the production of β-GlcNAc-phosphate-undecaprenol for subsequent transport to the outer leaflet and addition to the rhamnose chain, by two currently unknown enzymes (fig. 4).

The rhamnose backbone is a homogeneous structure composed of α-1,2 and α-1,3-linked residues which form an octameric repeating unit. It is therefore difficult to assign specific enzymes to each biosynthetic step, but unique structural features of the polymer can be extracted to propose a model for its biosynthesis. The first committed step of rhamnose backbone biosynthesis for example, must be the production of α-GlcNAc-pyrophosphoryl-undecaprenol (to act as an acceptor for rhamnose polymerisation), most likely catalysed by EpaA (supplementary Table 15). The following α-Rha residue (R_L_ in fig. 1g) is then most likely added by EpaB which shows the highest structural homology to *Mycobacterium tuberculosis* WbbL, an enzyme responsible for the production of α-Rha-(1,3)-α-GlcNAc-pyrophosphoryl-undecaprenol^34^. The *O*-methyl cap of the rhamnose backbone is also unique, and EpaN is the only candidate predicted to have methyltransferase activity. EpaN contains two other domains which are both predicted to have rhamnosyltransferase activity, suggesting that it is a processive enzyme able to catalyse the addition of the α-1,2 and α-1,3 linked rhamnose. Interestingly, these two enzymatic domains have high structural homology to EpaO, suggesting that EpaN and EpaO may have identical rhamnosyltransferase activity. An area of symmetry was observed within the rhamnose backbone (fig. 4, black box). The first three α-Rha residues of the octameric repeating unit (residues R_I_, R_F_ and R_B_, fig. 1f) are identical to the three terminal α-Rha residues (residues R_I_, R_C_ and R_A_, fig. 1g), and both substitute the previous α-Rha at the second position. We therefore propose that EpaO and EpaN recognise the same substrate and catalyse the addition of two α-1,2-Rha and an α-1,3-Rha. EpaN adds the final α-1,3-Rha and *O*-methylated it at the third position. EpaN therefore promotes the termination of α-rhamnose polymerisation since only one *O*-methyl cap is observed in the final structure (fig. 3b), and EpaO promotes polymerisation. We propose that the remaining rhamnosyltransferases within the EPA biosynthetic locus (EpaC and EpaD) are responsible for the addition of all other α-Rha residues although unknown rhamnosyltransferases outside of the EPA locus may also contribute towards rhamnose backbone biosynthesis.

Whilst the rhamnose backbone biosynthetic locus encodes an ABC transporter, the EPA decoration biosynthetic locus encodes a flippase (11709, fig. 1a) and an *O*-antigen ligase homolog (11710). This suggests that decoration building blocks are produced in the cytoplasm, but that polymerisation occurs after translocation to the outside of the cell (fig. 4).

The structures of the EPA mutants described in fig. 3a provide key insights into several of the step leading to the biosynthesis of EPA decorations. *EpaR* is a β-GalNAc priming glycosyltransferase, essential for EPA decoration biosynthesis (fig. 1b). It produces β-GalNAc-pyrophosphoryl-undecaprenol onto which the monomeric decoration subunit is built (fig. 4). The next step is the addition of a β-Glc to the nascent chain by the predicted β-1,3-glucosyltranferase 11720 (fig. 3a), (supplementary Table 16). 11715 then catalyses the addition of a β-Gal onto the third position of β-Glc (fig. 3a). The following addition of a glycerol-phosphate group onto the third position of β-Gal (fig. 4) could be catalysed either by 11711 or 11713, both of which are predicted glycerol-phosphate transferases and have highly similar predicted structures. Our structural data suggests that 11714 then catalyses the addition of β-GalNAc (fig. 3a), onto which another glycerol-phosphate is added, again by either 11711 or 11713. At this stage, we propose that the building block is translocated across the membrane. In analogy with the rhamnose backbone biosynthesis, we suggest that four additional modifications occur following exposure on the outer leaflet: (i) the addition of α-Glc residues onto the sixth position of β-GalNAc or (ii) onto the phosphoglycerol residue linked to β-GalNAc residues at the third position, (iii) the addition of α-GlcNAc onto β-Glc at the third position, and finally (iv) the addition of a phosphoglycerol onto this α-GlcNAc moiety at the sixth position. Our NMR data indicates that this α-GlcNAc-phosphoglycerol side chain (residue K/K* and Gro_4_, fig. 3c) is a major modification that requires the activity of 11706, an α-GlcNAc glycosyltransferase (supplementary Table 16 and fig. 3a).

The EPA rhamnose backbone and decorations are most likely ligated together into the final polymer by EpaP and/or EpaQ. Both proteins show high predicted structural homology to oligosaccharide transferases (supplementary Table 16). In agreement with this hypothesis, an epaQ mutation leads to a truncation of the EPA polymer^37^. We therefore suggest that EpaP/EpaQ binds to EPA decorations and catalyses the transfer of these subunits onto the rhamnose backbone. An unknown LytR–Cps2A–Psr (LCP) enzyme is likely responsible for the transfer of the complete EPA polymer onto the peptidoglycan layer for display at the surface^38,39^.

### GlcNAc-phosphoglycerol modifications contribute to host immune evasion

EPA decorations are essential for *E. faecalis* pathogenesis, protecting cells against phagocytosis by the host innate immune system^27^. The molecular basis of EPA decoration activity, however, remains unknown. We sought to use the epa decoration mutants described in this study to explore how specific structural determinants contribute towards innate immune evasion. We set up a phagocytosis assay using immortalised Bone Marrow-Derived Macrophages (iBMDMs) to measure the uptake of epa mutants constitutively expressing GFP^40^. As expected, the complete lack of decoration in the ΔepaR mutant (fig. 1g) led to a significant increase in phagocytosis (fig. 5a). We then tested epa mutants Δ*11720*, Δ*11715* and Δ*11714* which are all unable to produce polymerised EPA decorations (fig. 3a) and again found that these strains are more readily taken up by phagocytes (fig. 5b). Interestingly, polymerisation of EPA decoration subunits is not the only essential feature for immune evasion activity, since mutant Δ*11706* (lacking the GlcNAc-phosphoglycerol side chain substitution, fig. 3b) also exhibited a significantly reduced ability to evade phagocytosis (fig. 5c). These experiments therefore indicate that whilst EPA polymerisation is essential for *E. faecalis* phagocytosis evasion, specific EPA decoration motifs also play an important role during host-pathogen interactions.

### EPA recognition by bacteriophages

EPA decoration has been shown to be a major determinant of phage sensitivity for *E. faecalis*^22,41,42^. We therefore sought to determine the exact EPA decoration motifs required for the infection of *E. faecalis* OG1RF by probing the sensitivity of structurally diverse epa mutants against a collection of previously (Shef2^42^) and newly isolated (and uncharacterised) bacteriophages from local wastewater samples (OG_5, OG_6, OG_9, OG_14, OG_15 and OG_16).

We observed striking differences in the minimal structural motifs required for bacteriophage infectivity (fig. 6). Deletion of *11706*, which prevents the addition of the lateral α-GlcNAc-phospho-glycerol side chain, was sufficient to abolish infection by phage OG_16, whilst having a marginal effect on all other phage tested. Phage OG_9 appeared to require the terminal β-Gal-phosphoglycerol of EPA decoration, showing a 10-fold increase in virulence against Δ*11714* as compared to Δ*11715*. In contrast, the lytic activity of phages Shef2, OG_5, OG_14 and OG_15 was largely dependent on EPA decoration polymerisation, rather than a specific motif. A 100 to 1,000-fold decrease in plaque forming units was observed against Δ*11714* for Shef2, OG_5 and OG_14 respectively, and a complete lack of infection was recorded using OG_15. Interestingly phages OG_9 and OG_5 showed a low, residual efficiency of plating (ca. 1% and 0.01%, respectively) against mutant Δ*11715* and further truncations of EPA decorations had no impact, suggesting a role for other cell surface components during infection by these phages.

**Fig. 6.**
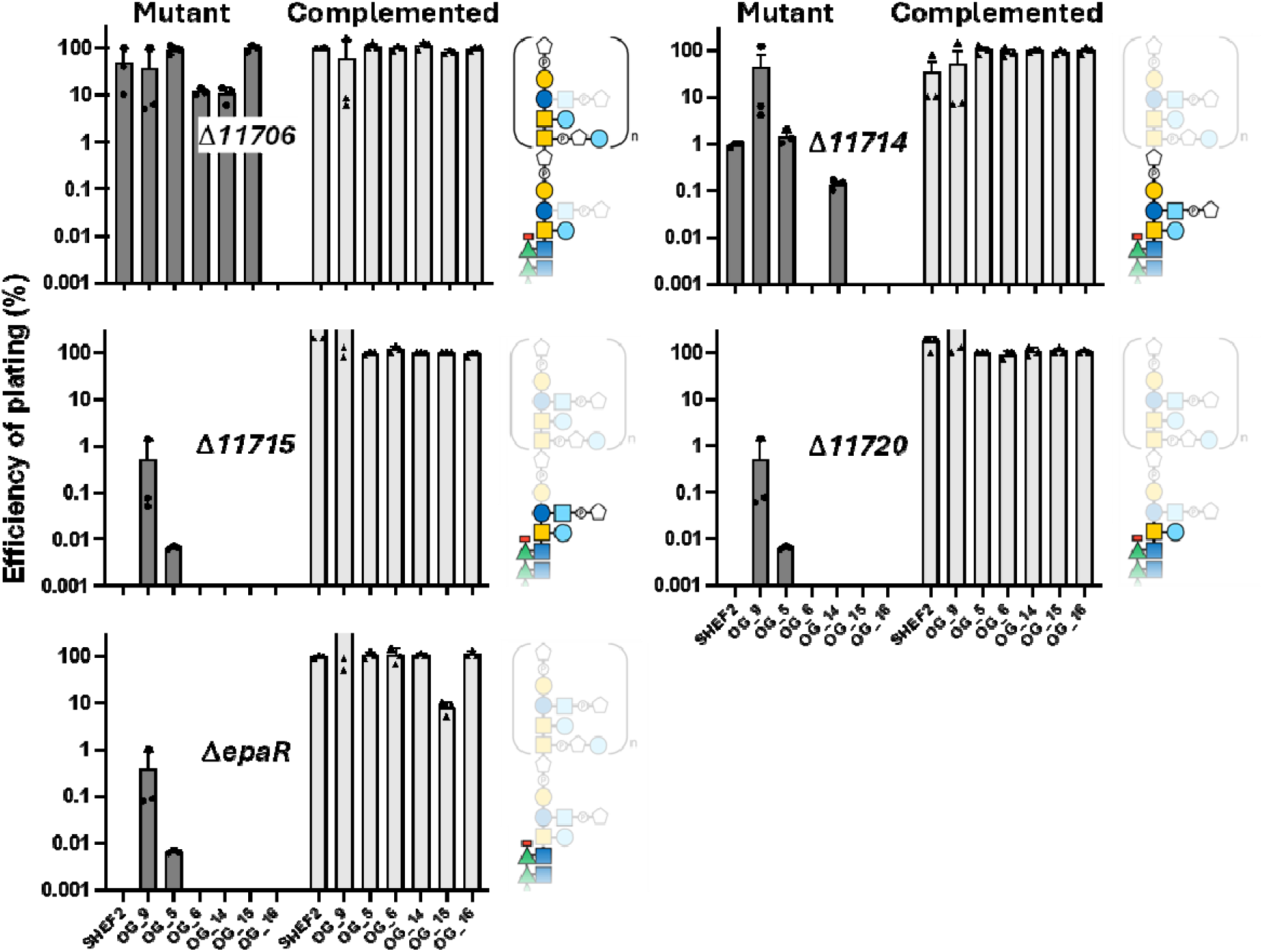
Efficiency of plating of bacteriophages targeting *E. faecalis* OG1RF against epa mutants. epa mutants were infected by six bacteriophages isolated using strain OG1RF as an indicator strain, OG_5, 6, 9, 14, 15, 16, and the previously described SHEF2^42^. The efficiency of plating was calculated by counting the number of plaque-forming units after serial dilutions of the phage stock. The number of plaques formed with OG1RF was used as a reference (100 %). The efficiency of plating is shown for individual mutants and their complemented derivatives. The structure of EPA decorations produced by each mutant is shown; the EPA motifs absent in each EPA structure can be seen by transparency.

## Discussion

EPA is a polymer conserved in Enterococci which plays a key role in growth, cell division, pathogenesis, and antimicrobial resistance^23^. Whilst the structure of EPA fragments has been described^28^, the complete structure of EPA has never been solved. Here, we used a repertoire of EPA biosynthetic mutants to elucidate the structure of the intact polymer by NMR and explored how EPA structure relates to its biological activity.

The high sequence identity between EpaA - EpaQ orthologs produced by *E. faecalis* strains^43^ suggests that the rhamnose backbone structure is conserved across *E. faecalis* strains. Unexpectedly, this work revealed differences between the EPA rhamnose backbone from E. faecalis OG1RF and the structure previously reported for V583^28^. The OG1RF rhamnose backbone repeating unit contains two extra α-Rha residues, one of which is substituted at the fourth position with an α-glucose. Whilst this modification has not been described in Lactococci or Enterococci^2^, it has recently been identified in *S. mutans*^44^. The glycosyltransferase responsible for catalysing the addition of this α-glucose is unknown and no homologs of *S. mutans* SccM and SccQ^44^ exist in *E. faecalis* OG1RF. The epa conserved region in V583 contains four genes with no counterpart in OG1RF, but none of these display sequence similarities with glycosyl transferases (supplementary fig. 6), therefore suggesting that unidentified glycosyltransferases located elsewhere on the chromosome may contribute to EPA backbone synthesis.

This work identified a set of four residues that cap the rhamnose backbone. The last rhamnose residue in this capping unit is *O*-methylated at the third position. In the E. coli O8 lipopolysaccharide mannan chain, this modification acts as a structural cue for transport across the membrane, thereby terminating polymerisation of the rhamnan chain^45,46^. We therefore propose that a similar mechanism is conserved in Enterococci. In agreement with this hypothesis, we confirmed the presence of an *O*-methyl group in V583 EPA, even though it was previously not reported (supplementary fig. 7). Since EpaN is the only enzyme conserved across Enterococci with predicted methyl transferase activity^43,47,48^ (supplementary Table 15), it is likely that this enzyme is adding the *O*-methyl cap.

EPA produced by the *epaR* mutant lacks decorations, supporting the idea that this gene is involved in the first committed step of EPA decoration biosynthesis. As *EpaR* is systematically present within the epa locus, and extremely conserved^43^, we propose that decorations are always built upon β-GalNAc-pyrophosphate-undecaprenol. Beyond the presence of a β-GalNAc residue in the repeating unit of decoration subunits, the composition and structure of EPA decorations are expected to reflect the high level of genetic diversity of the *epa* decoration loci. As anticipated, we therefore found that EPA decorations in OG1RF and V583 are radically different. OG1RF contains glycerol phosphate instead of ribitol phosphate, as well as more phosphodiester groups than V583 EPA which is also lacking any galactose residues. Distinct minor modifications were also observed within EPA decoration. These modifications are likely to occur after EPA is attached to the peptidoglycan layer as a dynamic response, or as a maturation process, like that observed in *Streptococcus*^9,49,50^.

Elucidating the structure of EPA produced by OG1RF and isogenic decoration mutants revealed the role of several enzymes in the biosynthesis of this polymer, allowing us to propose a biosynthetic pathway (fig. 4). Although some steps require further investigation, our model paves the way to fully characterise the contribution of individual EPA enzymes towards EPA biosynthesis. Our analysis suggests that some of the enzymes involved in EPA production, such as those catalysing the addition of α-Glc onto the backbone and/or decoration subunits, must be encoded outside of the epa biosynthetic cluster. Further work is needed to identify potential candidates.

Understanding the contribution of structural motifs to the biological activity of EPA requires access to EPA variants of known structures. A series of epa decoration mutants (Δ*11720*, Δ*11715*, Δ*11714* and Δ*11706*) allowed us to explore the mechanisms that underpin the evasion of phagocytosis and cell surface recognition by bacteriophages. As expected, the lack of polymerisation of EPA decorations (in mutants Δ*11720*, Δ*11715*, Δ*11714*) had a drastic impact on phagocytosis evasion (fig. 5). Nevertheless, the comparison between mutants Δ*11715* and Δ*11714* revealed that the terminal Gal-phosphoglycerol group (present in EPA_11714) plays an important role for the recognition by phagocytes. A striking result from the phagocytosis assay is the impact of the Δ*11706* mutation on phagocytosis. Despite the presence of polymerized decoration subunits in this mutant, the lack of the GlcNAc-phosphoglycerol side chain was sufficient to significantly impair evasion of phagocytosis. This result suggested that specific motifs play an important role in resistance against innate immunity.

The set of *epa* mutants was also used to explore the mechanisms underpinning OG1RF lysis by bacteriophages (fig. 6). This work revealed an unexpected diversity of cell surface recognition strategies by bacteriophages and provided evidence that EPA is not the only determinant of binding. The proof of concept provided in this work is an important step towards identifying distinct classes of bacteriophages that target different surface structures for the design of phage cocktails as therapeutics against these pathogens. Furthermore, the combination of phage sequencing, with the molecular characterisation of the EPA motif required for phage virulence will provide information about receptors proteins to engineer phages with tailored host ranges.

The cell envelope composition in *E. faecalis* is complex. Whilst all strains produce EPA, some (such as V583) also produce a facultative capsular polysaccharide^51^ absent in OG1RF^52^. Why some strains produce a capsule and others do not is currently unclear. Additionally, in *E. faecalis* V583, a third polymer of low molecular weight has also been described and proposed to be a teichoic acid polymer^51^. The genetic and structural characterization of this polymer awaits further analysis. Both OG1RF and V583 represent ideal models which can be used as platform strains to explore the distinct activities of cell wall polysaccharides and the interplay between the capsule, EPA and putative wall teichoic acids on the *E. faecalis* cell surface. In this work, we made the first steps towards the complete molecular characterisation of EPA using isogenic deletion mutants. Future work may also take advantage of heterologous expression strategies previously described^40,53^ to enable the comparison of isogenic strains and explore strain-strain variability. Elucidating the structure/function relationship of EPA will enable the development of vaccines and novel phage therapies for the prevention and treatment of enterococcal infections.

## Methods

### Bacterial strains, plasmids, oligonucleotides, and growth conditions

Bacterial strains, plasmids and oligonucleotides used in this study are described in supplementary Table 17. *E. coli* TG1 was grown at 37°C in Luria-Bertani supplemented with 200 µg/mL erythromycin. *E. faecalis* was grown in Brain Heart Infusion at 37°C without agitation. The media was eventually supplemented with 30 µg/mL erythromycin and 10 ng/mL of anhydrotetracycline to maintain pTet derivatives and induce the epa gene expression they encode, respectively. Tetracycline was added at 5µg/mL to maintain the pMV158-GFP plasmid.

### Construction of the OG1RF_*11706* mutant

Four mutants used in this study were previously described^22^, ΔOG1RF_*11714*, ΔOG1RF_*11715* and ΔOG1RF_*11720* ^27^). Mutant ΔOG1RF_*11706* was built by allelic exchange using the strategy previously described^54^ (Two homology regions flanking the OG1RF_*11706* open reading frame were amplified from genomic DNA via PCR: a 5’ homology region, using primers SM_0216 (sense) and SM_0217 (antisense) and a 3’ homology region, using primers SM_0218 (sense) and SM_0219 (antisense). Once purified, the two PCR products were mixed in equimolar amount and fused via splice overlap extension PCR^55^ using primers SM_0216 and SM_0219. The resulting fragment was cut by XhoI and NotI and cloned into pGhost9^56^. Candidate pGhost derivatives were screened by PCR using primers SM_0171 and SM_0172. A positive clone containing the fused H1-H2 insert was checked by Sanger sequencing and the corresponding plasmid was named pGhost-11706.

pGhost-11706 was electroporated into *E. faecalis* OG1RF. Transformants were selected at 28°C and streaked onto BHI agar containing 30 μg/mL erythromycin at 42°C (a non-replication-permissive temperature) to select single crossover recombination events. One colony was used to make subcultures at 28°C without antibiotic. To find double crossover recombination events, single colonies were re-isolated and screened via PCR using the primers SM_0256 and SM_0257.

For complementation, the complete OG1RF_*11706* open reading frame was PCR amplified using primers SM_0348 and SM_0349 and cloned into the pTetH vector using SacI and BamHI. Candidates were screened by using primers SM_0100 and SM_0101. A positive clone was checked by Sanger sequencing and named pTet-11706.

### EPA purification

EPA was purified from 1.8 L cultures. Cell walls were first purified by boiling the cells in 5% (w/v) SDS for30 minutes. After 5 washes in MilliQ water, cell walls were digested overnight at 37°C with 5,000 U of mutanolysin in 10 mM sodium phosphate buffer (pH 5.5) in a volume of 5 mL. Soluble cell wall fragments were injected on a Superdex200 26/60 gel filtration column equilibrated in MilliQ water. Fractions containing neutral sugars were identified either using a phenol-sulphuric acid assay^57^ or by absorbance at 206 nm and pooled. A further anion exchange chromatography step was carried out on a 5 mL DEAE FF column equilibrated in MilliQ water. EPA was eluted with a linear gradient to 500 mM NaCl. The fractions containing EPA were detected using a neutral sugar assay or by absorbance at 206 nm, pooled, concentrated to 2 mL and injected on a HiPrep 26/10 desalting column. The final, desalted EPA fractions were freeze-dried for NMR analysis.

### Acid hydrolysis of EPA

To produce EPA fragments OS1 and OS2, 3 mg of purified EPA were treated with 48% (v/v) hydrofluoric acid and incubated at 4 °C for 18 hours. Hydrolysis products were separated using a self-packed column of P-4 fine resin (Bio-Rad) and identified using the neutral sugar assay. The OS3 fragment was produced using 28 mg of purified EPA treated with 48% (v/v) hydrofluoric acid for 1 hour at room temperature. The EPA partial cleavage products were then desalted using Sephadex G15 and OS3 separated by reversed-phase HPLC on a C18 hypersil column using a water - methanol gradient in the presence of 0.05 % (v/v) TFA.

### NMR experiments

NMR spectra were recorded on a 900 MHz Bruker Advance NEO using a 5-mm-diameter triple-resonance inverse cryoprobe (^1^H and ^13^C; EPA fragments OS1 and OS2), a 600 MHz Bruker AVANCE III using a 5 mm PABBO BB-1H/D Z-GRD Probe (^1^H,^13^C, and ^31^P; OS3), a 600 MHz NEO using a 5-mm-diameter TCI cryo-probe (^1^H and ^13^C; EPA, EPA_epaR, EPA_11720, EPA_11715, EPA_11714 and EPA_11706) and a 500 MHz Bruker Advance II using a 5-mm-diameter BBO probe (^1^H and ^31^P; OS1, OS2, EPA, EPA_epaR, EPA_11720, EPA_11715, EPA_11714 and EPA_11706). All experiments were performed at 298 K in D_2_O. Samples were supplemented with 0.01 % (v/v) acetone as a reference to calibrate ^1^H and ^13^C chemical shifts which are reported as parts per million (δ_H_ 2.225 ppm and δ_C_ 31.55 ppm).

^1^H-^1^H COSY NMR experiments used 2D homonuclear shift correlation using gradient pulses for selection and presaturation the during relaxation delay. ^1^H-^1^H-TOCSY NMR experiments were recorded using mixing times of 80 and 120 ms, and ^1^H-^1^H-NOESY NMR experiments were recorded using mixing times of 50 ms. ^1^H-^1^H-ROESY NMR experiments were phase-sensitive using Echo/Antiecho-TPPI gradient selection. A phase-sensitive multiplicity-edited sequence was used to record the ^1^H-^13^C-HSQC which used echo/anti-echo-TPPI gradient selections with decoupling during acquisition and trim pulses in the Inept transfer. The ^1^H-^13^C-HSQC-TOCSY NMR experiments were acquired using MLEV17 for homonuclear Hartman-Hahn mixing and echo/anti-echo-TPPI gradient selections with decoupling during acquisition and trim pulses in the Inept transfer, and a mixing time of 100 ms. The ^1^H-^13^C-HMBC NMR experiments were optimised on long-range couplings ^n^J_CH_ of 8 Hz without multiplicity selection or decoupling during acquisition. The ^1^H-^31^P-HSQC-TOCSY NMR experiments were phase sensitive and acquired using MLEV17 for homonuclear Hartman-Hahn mixing with decoupling during acquisition and trim pulses in the Inept transfer and a mixing time of 30 ms. The ^1^H-^31^P-HSQC-TOCSY NMR experiment was phase-sensitive and acquired using MLEV17 for homonuclear Hartman-Hahn mixing with decoupling during acquisition with a mixing time of 100 ms. Spectra were processed on Topspin 3.2 and analysed on Topspin 4.0.9.

### Phagocytosis susceptibility assay

Immortalised bone marrow-derived macrophages (iBMDMs) from wild type mice were obtained from the BEI Resources, NIAID NIH (NR-9456) (https://www.beiresources.org/Catalog/cellBanks/NR-9456.aspx)^57^. iBMDMs were cultured in DMEM (Gibco) supplemented with 1% (v/v) foetal bovine serum (FBS, PAN Biotech; low endotoxin, heat-inactivated), penicillin (10 U/mL)/streptomycin (1 mg/mL) and 1 mM sodium pyruvate. Cells were cultured in standard tissue culture flasks or multi-well plates at 37°C in 5% CO_2_, washed in PBS and given fresh media once every 48 hours. On day 1, iBMDMs were checked for >70% confluence and diluted in fresh media to a concentration of 5 x 10^5^ cells per well. *E. faecalis* overnight cultures were set up. On day 2, *E. faecalis* cultures were started (100 μl overnight into 10 mL fresh media) and grown at 37°C until OD_600_ ≈ 0.3. Cultures were pelleted (5 min at 4000 x *g*) and resuspended in an equal volume of DMEM (serum- and antibiotic-free). iBMDMs were washed and given fresh media and supplemented with 2.5 x 10^6^ CFU bacteria per well (MOI = 5). Three wells were allocated per bacterial strain in each experiment. After 1 hour at 37°C in 5% CO_2_, iBMDMs were washed and treated with 250 μg/mL gentamycin + 20 μg/mL vancomycin for 1 hour at 37°C in 5% CO_2_. Cells were then washed twice with PBS and detached by treating with 1 mL Accutase™ (Merck) for 30 min at 37°C in 5% CO_2_. Detached iBMDMs were pelleted (5 min at 7,000 x g), fixed, resuspended in 200 μl filtered PBS, and stored at 4°C in the dark.

For flow cytometry analysis, 200 μl of each iBMDM sample were transferred to a 96-well plate. Data acquisition was performed using a Guava easyCyte HT flow cytometer (Luminex). Data analysis was carried out using guavaSoft v 3.1.1; gating strategy is shown in Supplementary fig. 8.

### Bacteriophage infections

Phages were isolated using the agar overlay method as previously described^42^. (*E. faecalis* indicator strains were grown to exponential phase (OD_600_ ≈ 0.5) in BHI supplemented with 5 mM MgSO_4_ and 5 mM CaCl_2_ and mixed with and equal volume of wastewater. After 10 min at room temperature, 1mL of the mixture was added to 4 mL of BHI-top agar (0.6% w/v) and immediately spread on a BHI-agar plate. Individual plaques were purified through two additional rounds of infections. Pure plaques were resuspended in 200 µL of BHI, gently vortexed for 10 min and used to prepare plate lysates using the overlay method previously described^42^. Plates with confluent plaques were used to prepare high-titre stocks. Three mL of SM buffer (50 mM Tris-HCl pH7. 5, 100 mM NaCl, 5m M MgSO_4_) were added on top of plates, left under agitation for 3 hours and the suspension was filtered through a Millipore membrane (0.45 µm). Efficiency of plating was determined using the method described above using serial dilutions of phage stocks (10^6^-10^9^ pfu/mL). The number of pfu/mL measured with OG1RF as an indicator strain was used to define 100% infection.

### Statistical analysis

GraphPad Prism version 10.1.2 was utilised for statistical analysis. Error bars on graphs represent mean ± SD. Each set of iBMDM flow cytometry data was analysed using a one-way ANOVA with Welch’s correction, followed by Dunnett’s multiple comparisons test.

## Acknowledgements

JLD and RES were funded by the White Rose Doctoral Training Programme (BBSRC grant BB/ M011151/1). JLD and RES were awarded a MOBLILEX scholarship from Université de Lille, France and a short-term EMBO Fellowship, respectively for specific training in polysaccharide analysis in YG’s laboratory. JLD was also supported by the Publication Scholarship scheme funded by the University of Sheffield. JSN was supported by a studentship from the DiMeN Doctoral Training Programme (Medical Research Council grant MR/N013840/1). We thank Martin Stranex (Sheaf and Porter Rivers Trust) and Oliver Waite (Blackburn Meadows wastewater treatment plant) for their contribution to the collection of wastewater samples for phage isolation. Dr Kristi Frank (Uniformed Services University of the Health Sciences, Bethesda) is acknowledged for the kind gift of the *epaR* mutant and its complemented derivative. We would also like to acknowledge the specialist support from the Plateforme d’Analyse des Glycoconjugués (PAGés, Faculté des Sciences et Technologies de Lille). An upgrade to the 600 MHz NMR spectrometer was funded by the Biotechnology and Biological Sciences Research Council (BB/R000727/1).

