## Supplementary Figures for "The Enterococcal Polysaccharide Antigen: from structure to biosynthesis and function"

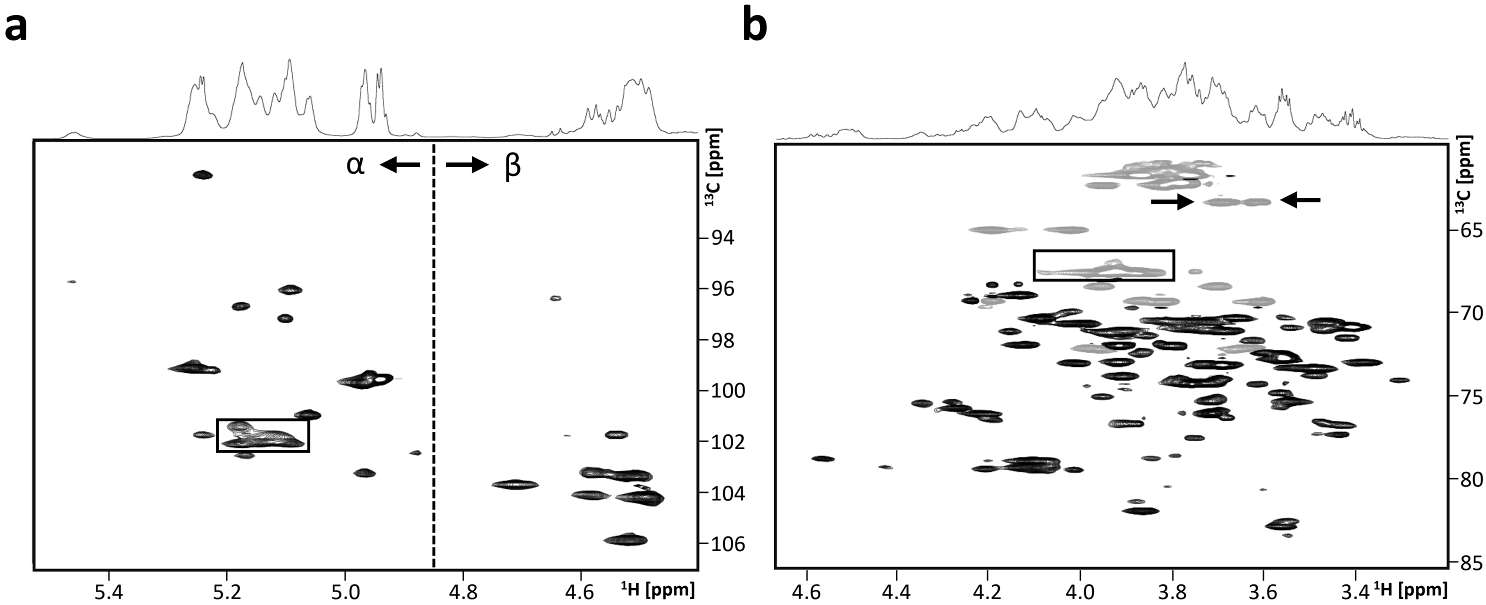


**Supplementary Fig. 1. 2D NMR analysis of *E. faecalis* OG1RF EPA by $^{1}$H-$^{13}$C-HSQC.**

**a**, Anomeric region. Each peak corresponds to an anomeric $^{13}$C and $^{1}$H pair of an EPA monosaccharide residue. The large signal between 5.07-5.2/101.2-102.2 ppm (boxed) corresponds to multiple peaks from monosaccharides of similar configuration and chemical environment. Monosaccharides with an anomeric $^{1}$H shift of above 4.85 ppm (dashed line) correspond to α-sugars, whilst those below 4.85 ppm correspond to β-sugars. **b**, Carbohydrate ring system region. $CH_2$ groups are shown in the negative phase (grey). Glycerol was detected within EPA, with characteristic shifts at 3.6-3.7/63.4 ppm corresponding to the $CH_2OD$ group of unsubstituted glycerol (arrows), and 3.8-4.0/67.6 ppm (boxed) corresponding to the $CH_2PO_4X$ group of glycerol when phosphorylated (where X detonates an unknown chemical group).

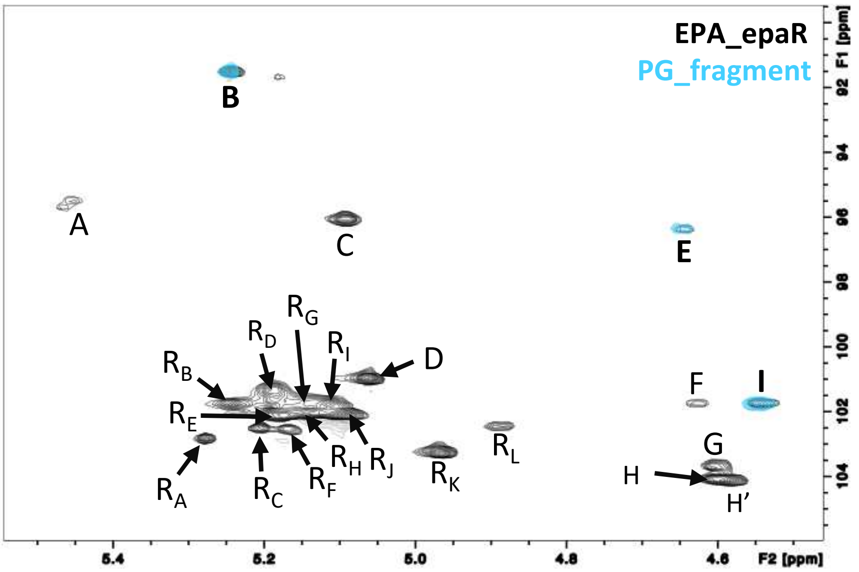


**Supplementary Fig. 2. Identification of $^{1}H$-$^{13}C$ signals corresponding to the peptidoglycan moiety within EPA_epaR.**

Across this study, all EPA samples were purified enzymatically with mutanolysin, leaving a monomeric peptidoglycan scar which contains a MurNAc residue existing in both an α and β form. To identify the anomeric $^{1}H$-$^{13}C$-HSQC signals produced by these peptidoglycan fragments, cell wall sacculi was purified from *E. faecalis* OG1RF and digested with mutanolysin. Monomeric peptidoglycan fragments were purified by size exclusion chromatography and a $^{1}H$-$^{13}C$-HSQC 2D NMR experiment was recorded (blue). The anomeric region was overlayed with the $^{1}H$-$^{13}C$-HSQC spectrum of EPA_epaR sample (black). Residues B, E and I (bold) have identical shifts in both samples and must therefore correspond to the three anomeric signals (α-MurNAc, β-MurNAc and β-GlcNAc) from the peptidoglycan scar.

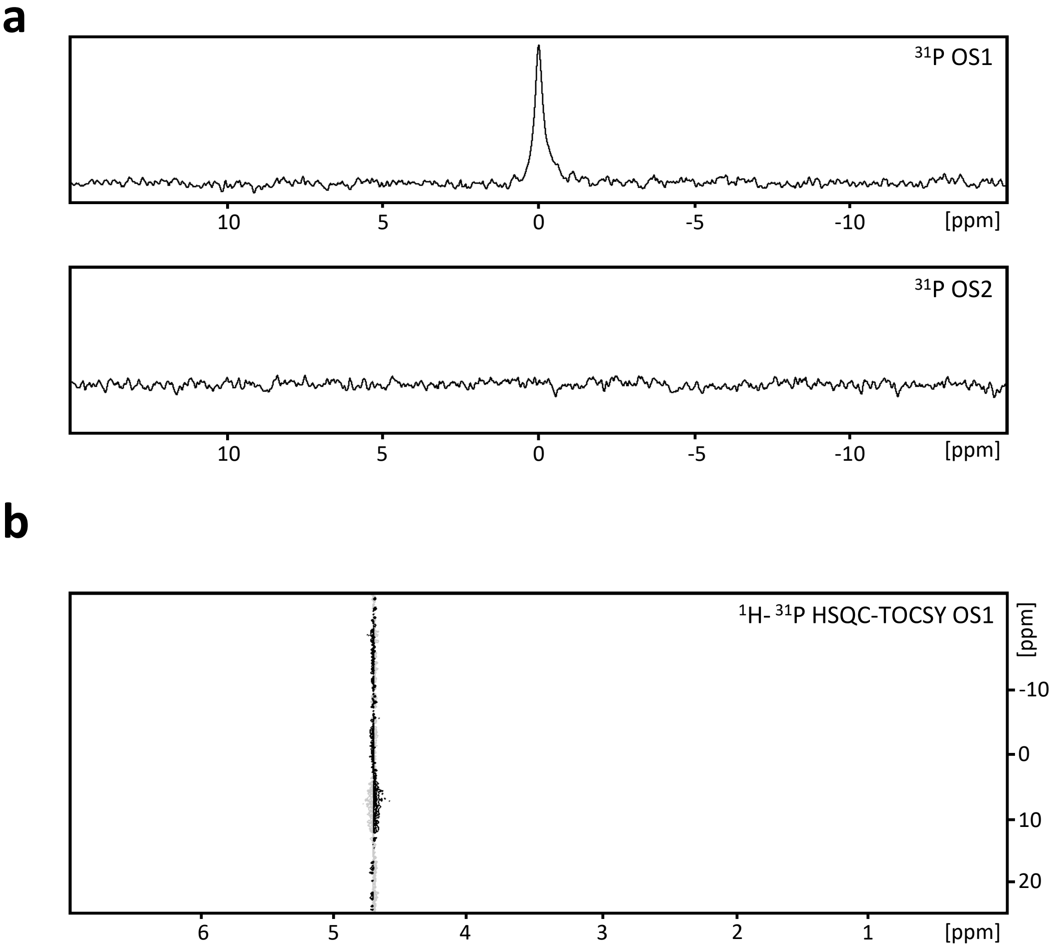


**Supplementary Fig. 3 Characterization of phosphorus signals within OS1 and OS2. a,** $^{31}P$ 1D NMR experiment was recorded on OS1 (top) and OS2 (bottom) fragments. Only one signal at 0.0 ppm was observed across the two samples, in OS1, most likely corresponding to a free phosphate group ($D_3PO_4$). **b,** To confirm this a $^{1}H$- $^{31}P$-HSQC-TOCSY NMR experiment was recorded on OS1, and as expected, no cross peaks were detected.

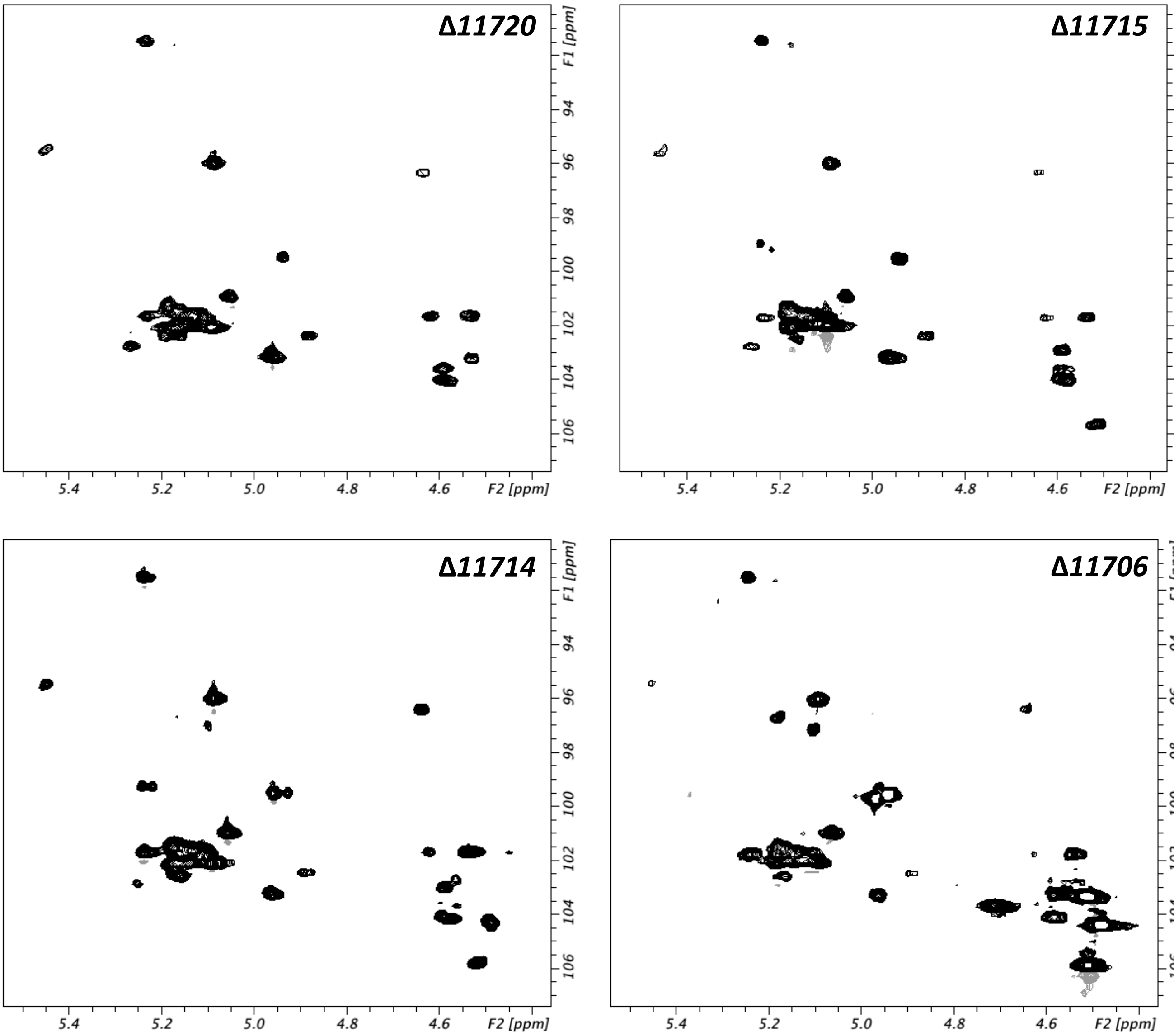


**Supplementary Fig. 4. Initial analysis of EPA produced by *epa* glycosyltransferase mutants.** The anomeric regions of $^{1}H$-$^{13}C$-HSQC spectra recorded on EPA purified from *epa* mutants (*Δ11720* top left, *Δ11715* top right, *Δ11714* bottom left and *Δ11706* bottom right).

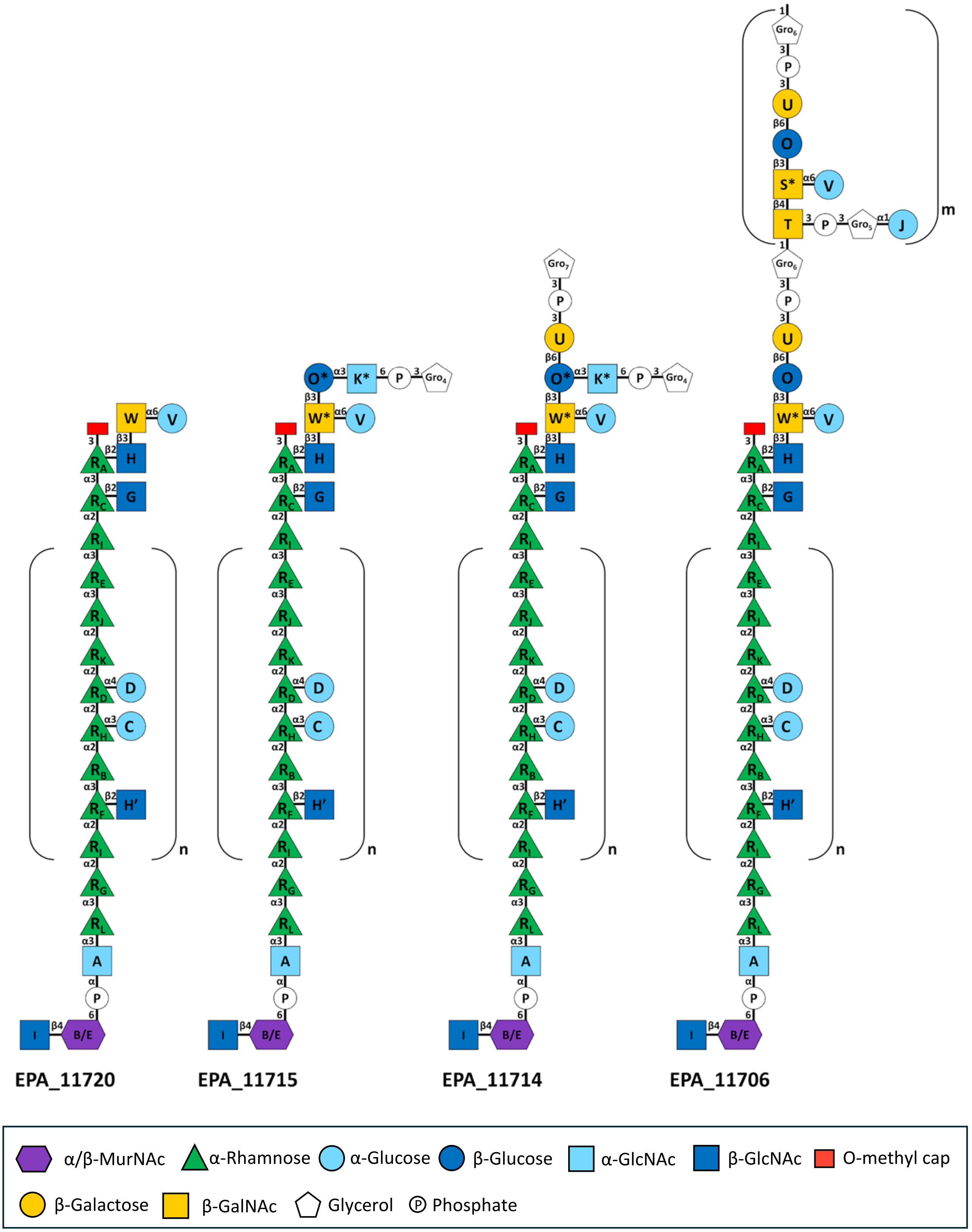


**Supplementary fig. 5. Full structures (from left to right) of EPA_11720, EPA_11715, EPA_11714 and EPA_11706.** Residues are labelled according to supplementary Tables 1, 7, 9, 11 and 13.

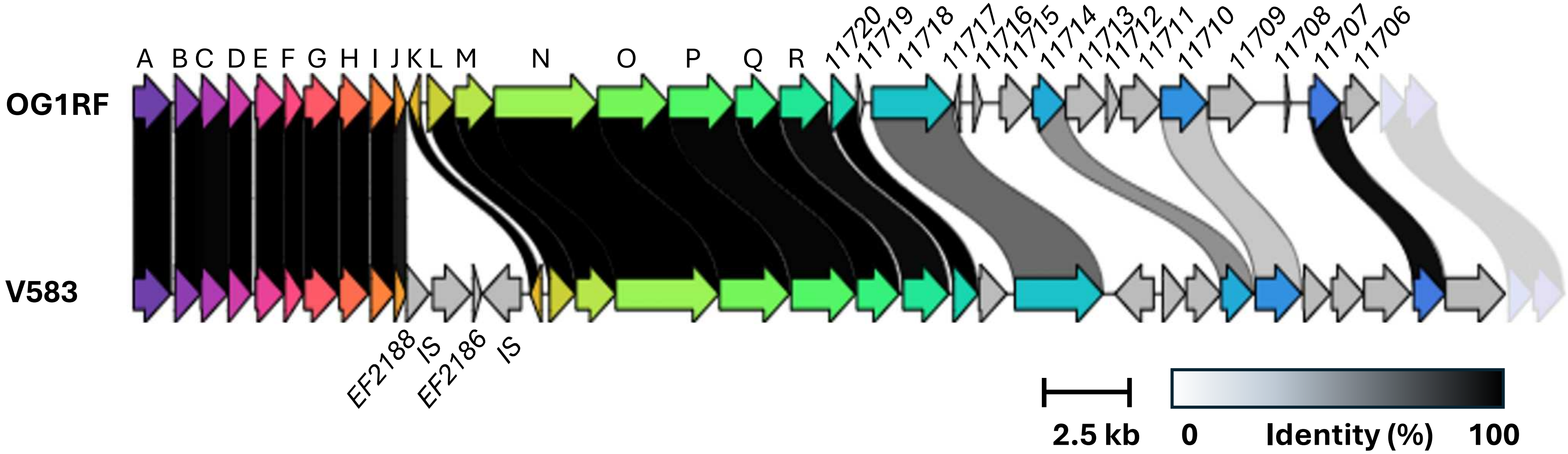


**Supplementary figure 6. Sequence comparison of OG1RF and V583 *epa* loci.** The conserved region (*epaA*-*epaR*) encodes 18 proteins with >95% sequence identity. Strain V583 contains 4 genes with no counterpart in OG1RF and no predicted function associated with carbohydrate metabolism: EF2188, and EF2186 share homology with a putative bifunctional racemase/acetyltransferase but correspond to a truncated gene resulting from the insertion of *EF2187* and *EF2185* encoding two IS transposases. Very few genes in the variable region are conserved, apart from *11720* encoding a putative α-1,3 glucosyltransferase; *11718*, a peptidoglycan hydrolase hydrolase; *11714*, encoding a β-1,3-galactosyltransferase, *11710*, encoding a ligase, *11707*, encoding an UDP-glucose 4-epimerase.

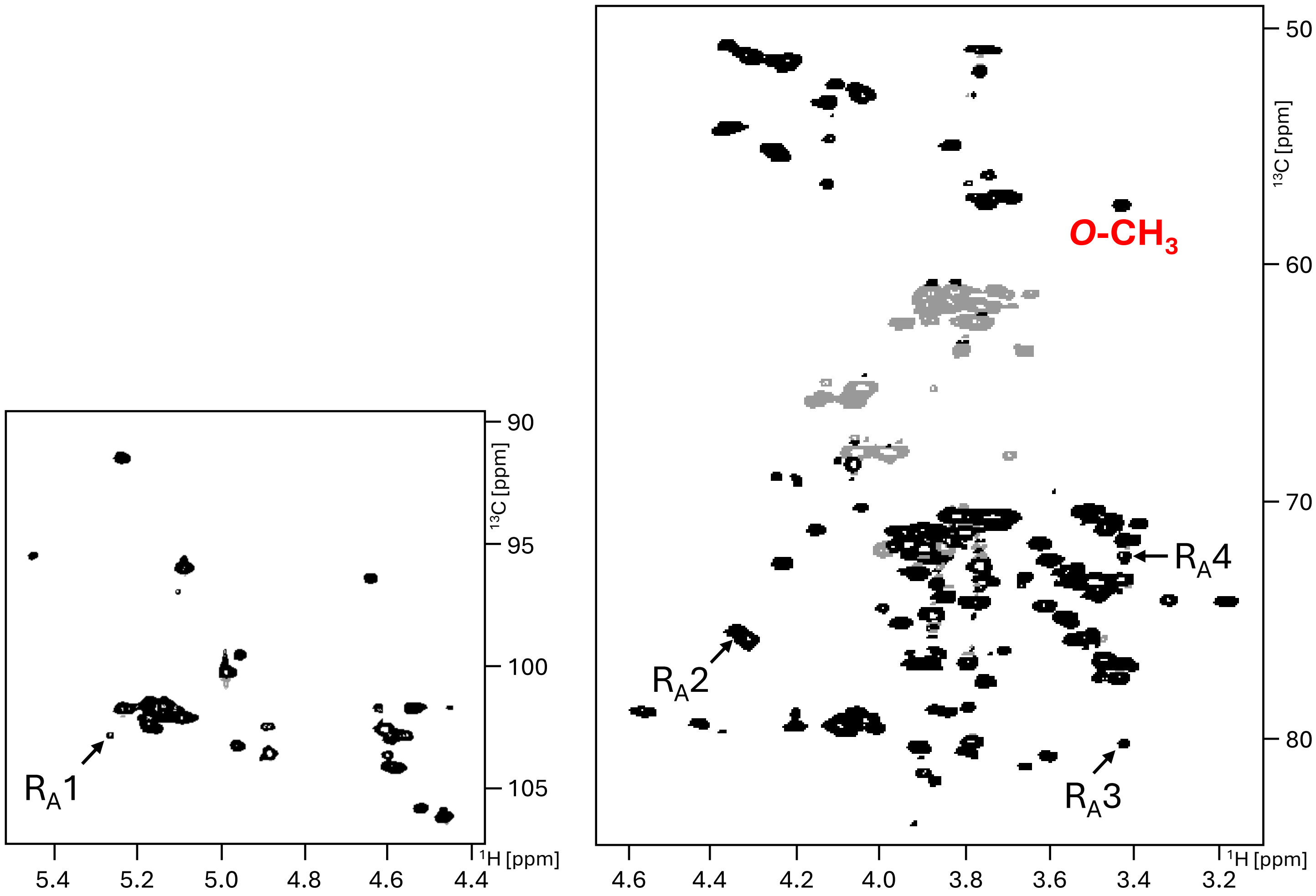


**Supplementary fig. 7. The rhamnose *O*-methyl cap is also present in EPA purified from *E. faecalis* V583.**

Closer analysis of an $^{1}H$-$^{13}C$ HSQC recorded on EPA purified from *E. faecalis* V583 shows that rhamnose residue $R_A$ (see Fig. 1g), which is methylated at the third position and terminates rhamnose backbone polymerisation in OG1RF, is present in EPA from V583. The anomeric shift (left) of rhamnose residue $R_A$ and shifts corresponding to $R_A$ positions 2-4 and the *O*-methyl group substituting this residue at position 3 (right), are all identical to those observed in EPA purified from *E. faecalis* OG1RF.

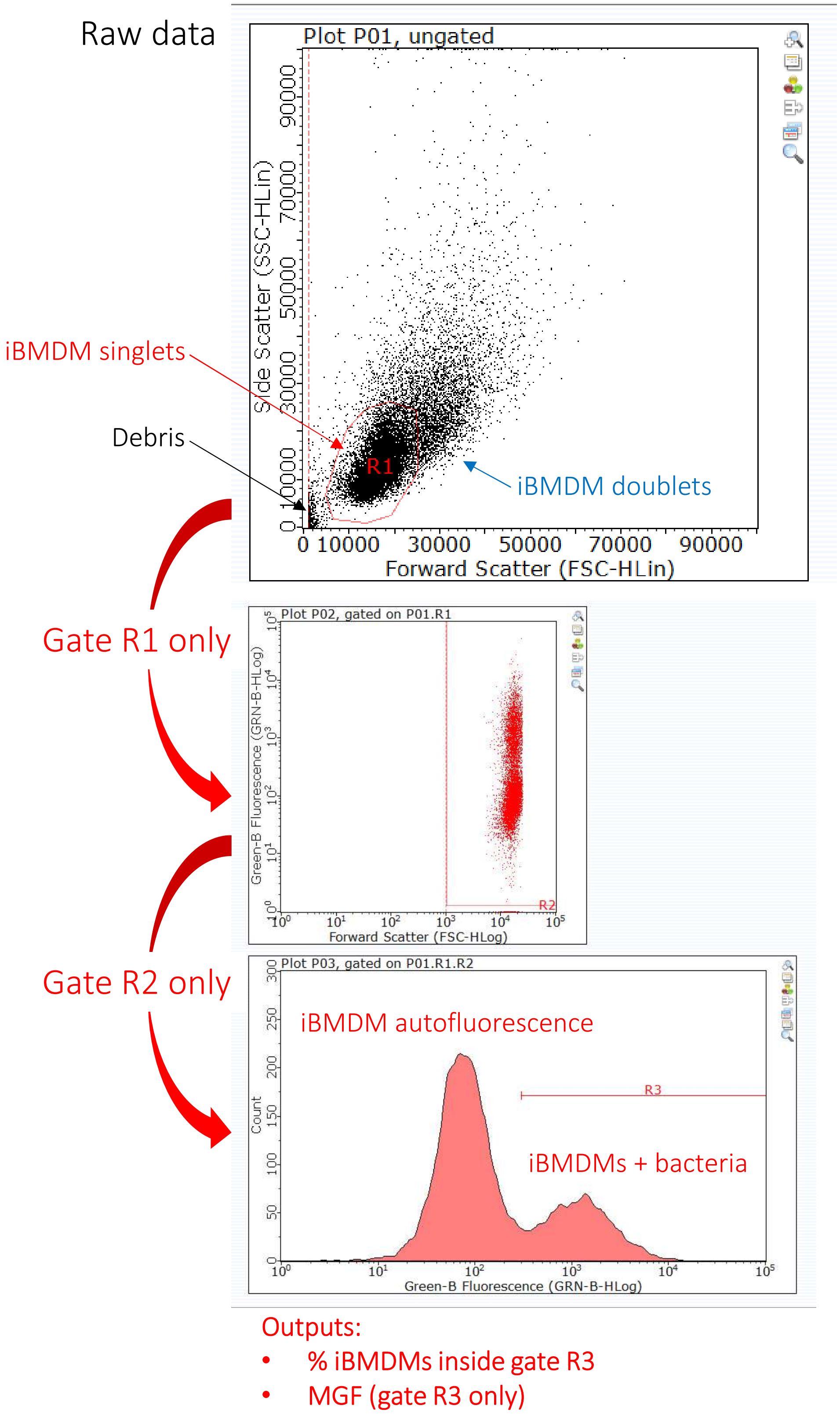


**Supplementary Fig. 8: Gating strategy for flow cytometry analysis of iBMDMs using GuavaSoft 3.1.1.** Debris and cell clumps were excluded from gate R1. Gate R1 data was re-plotted as FSC log (x axis) versus green fluorescence log (y axis). Gate R2 excluded more debris. Gate R2 data was plotted as a histogram (green fluorescence log (x axis) versus count (y axis)). The left peak (peak green fluorescence ≈ 7 x $10^1$) corresponds to autofluorescence of empty iBMDMs, whereas the right peak corresponds to iBMDMs with internalised GFP-labelled bacteria. Gate R3 (green fluorescence > 3 x $10^2$) was drawn to select only the right peak. The percentage of iBMDMs containing bacteria was calculated using (no. iBMDMs in gate R3/no. iBMDMs in gate R2) x 100. The median green fluorescence (MGF) of gate R3 data was calculated by GuavaSoft 3.1.1.
