## Supplementary Tables for "The Enterococcal Polysaccharide Antigen: from structure to biosynthesis and function"

**Supplementary Table 1.** All $^{1}H$, $^{13}C$ and $^{31}P$ NMR chemical shifts (ppm) at 298 K of EPA purified from Δ*epaR* cells (EPA_epaR; Ho *et al.*, 2018) determined through the analysis of $^{1}H$-$^{1}H$-COSY, $^{1}H$-$^{1}H$-TOCSY, $^{1}H$-$^{13}C$-HSQC, $^{1}H$-$^{13}C$-HSQC-TOCSY, $^{31}P$ and $^{1}H$-$^{31}P$-HSQC-TOCSY experiments.

| **Residue** | | ***H1***<br>**C1** | ***H2***<br>**C2** | ***H3***<br>**C3** | ***H4***<br>**C4** | ***H5***<br>**C5** | ***H6***<br>**C6** | **H6'**<br>**C6** | **N-acetyl**<br>**$CH_3$** | ***O*-methyl**<br>**$CH_3$** | **$PO_4$** |
|---|---|---|---|---|---|---|---|---|---|---|---|
| →3)-α-GlcNAc*p*-(1-*P* | **A** | *5.46*<br>95.6 | *4.12*<br>54.6 | *3.81*<br>80.5 | *3.60*<br>69.5 | *3.90*<br>74.7 | *ND*<br>*ND* | *ND*<br>*ND* | *2.09*<br>23.5 | | -1.79 |
| →4)-*P*-6-α-MurNAc*p*-OH | **B** | *5.24*<br>91.5 | *3.83*<br>54.9 | *3.76*<br>77.5 | *3.89*<br>76.7 | *3.87*<br>72.3 | *4.07*<br>64.9 | *4.13*<br>64.9 | *2.08*<br>23.7 | | -1.79 |
| α-Glc*p*-(1→ | **C** | *5.09*<br>96.0 | *3.60*<br>72.3 | *3.78*<br>74.2 | *3.46*<br>70.7 | *3.91*<br>72.9 | *3.83*<br>61.7 | *ND*<br>61.7 | | | |
| α-Glc*p*-(1→ | **D** | *5.06*<br>101.0 | *3.56*<br>72.9 | *3.70*<br>74.0 | *3.46*<br>70.5 | *4.01*<br>73.1 | *3.81*<br>61.4 | *ND*<br>61.4 | | | |
| →4)-*P*-6- β-MurNAc*p*-OH | **E** | *4.64*<br>96.4 | *3.78*<br>57.1 | *3.60*<br>80.7 | *3.86*<br>*76.5* | *3.48*<br>76.5 | *4.13*<br>64.9 | *4.19*<br>64.9 | *2.08*<br>23.7 | | -1.79 |
| β-GlcNAc*p*-(1→ | **F** | *4.63*<br>101.7 | *3.79*<br>57.1 | *3.55*<br>75.2 | *3.42*<br>71.5 | *3.48*<br>77.3 | *ND*<br>*ND* | *3.96*<br>62.5 | *2.08*<br>23.7 | | |
| β-GlcNAc*p*-(1→ | **G** | *4.60*<br>103.7 | *3.69*<br>57.0 | *3.61*<br>74.3 | *3.47*<br>71.0 | *3.42*<br>76.9 | *3.77*<br>61.7 | *3.88*<br>61.7 | *2.08*<br>23.7 | | |
| β-GlcNAc*p*-(1→ | **H** | *4.61*<br>104.0 | *3.71*<br>57.0 | *3.58*<br>74.7 | *3.47*<br>71.0 | *3.42*<br>76.90 | *3.76*<br>61.7 | *ND*<br>61.7 | *2.08*<br>23.7 | | |
| β-GlcNAc*p*-(1→ | **H'** | *4.59*<br>104.1 | *3.70*<br>57.0 | *3.61*<br>74.3 | *3.47*<br>71.0 | *3.42*<br>76.9 | *3.87*<br>61.7 | *ND*<br>61.7 | *2.08*<br>23.7 | | |
| β-GlcNAc*p*-(1→ | **I** | *4.54*<br>101.7 | *3.75*<br>57.3 | *3.57*<br>74.8 | *3.42*<br>71.5 | *3.44*<br>77.3 | *3.77*<br>62.4 | *3.95*<br>62.4 | *2.08*<br>23.7 | | |
| →2)-3-*O*-methyl-α-Rha*p*-(1→ | **$R_A$** | *5.28*<br>102.8 | *4.34*<br>75.9 | *3.44*<br>80.2 | *3.42*<br>72.4 | *3.73*<br>70.6 | *1.32*<br>18.1 | | | *3.44*<br>57.4 | |
| →2)-α-Rha*p*-(1→ | **$R_B$** | *5.24*<br>101.8 | *4.01*<br>79.5 | *3.86*<br>71.4 | *3.54*<br>73.3 | *3.73*<br>70.9 | *1.34*<br>18.1 | | | | |
| →2,3)-α-Rha*p*-(1→ | **$R_C$** | *5.20*<br>102.5 | *4.21*<br>78.9 | *3.91*<br>78.8 | *ND*<br>73.9 | *3.72*<br>70.9 | *1.28*<br>17.9 | | | | |
| →2,4)-α-Rha*p*-(1→ | **$R_D$** | *5.19*<br>101.3 | *4.06*<br>78.8 | *4.03*<br>70.0 | *3.55*<br>82.5 | *3.88*<br>69.9 | *1.38*<br>18.2 | | | | |
| →2)-α-Rha*p*-(1→ | **$R_E$** | *5.18*<br>102.2 | *4.07*<br>79.2 | *3.95*<br>71.3 | *3.50*<br>73.4 | *3.83*<br>70.6 | *1.32*<br>18.1 | | | | |
| →2,3)-α-Rha*p*-(1→ | **$R_F$** | *5.17*<br>102.5 | *4.21*<br>79.4 | *3.89*<br>76.8 | *ND*<br>73.9 | *3.72*<br>70.9 | *1.28*<br>17.9 | | | | |
| →2)-α-Rha*p*-(1→ | **$R_G$** | *5.16*<br>102.0 | *4.07*<br>79.1 | *3.95*<br>71.3 | *3.50*<br>73.4 | *3.83*<br>70.6 | *1.32*<br>18.1 | | | | |
| →2,3)-α-Rha*p*-(1→ | **$R_H$** | *5.15*<br>102.1 | *4.34*<br>75.5 | *3.95*<br>75.2 | *3.63*<br>71.7 | *3.74*<br>70.7 | *1.31*<br>18.1 | | | | |

| | | | | | | | |
|---|---|---|---|---|---|---|---|
| →2)-α-Rha*p*-(1→ | **R$_I$** | *5.11*<br>101.7 | *4.13*<br>79.2 | *3.91*<br>71.2 | *3.47*<br>73.4 | *3.81*<br>70.5 | *1.32*<br>18.1 |
| →2)-α-Rha*p*-(1→ | **R$_J$** | *5.10*<br>102.0 | *4.10*<br>79.3 | *3.90*<br>71.2 | *3.49*<br>73.3 | *3.74*<br>70.7 | *1.29*<br>17.9 |
| →3)-α-Rha*p*-(1→ | **R$_K$** | *4.96*<br>103.2 | *4.16*<br>71.1 | *3.85*<br>78.8 | *3.56*<br>72.9 | *3.77*<br>70.6 | *1.29*<br>17.9 |
| →3)-α-Rha*p*-(1→ | **R$_L$** | *4.89*<br>102.4 | *3.83*<br>71.8 | *3.80*<br>78.6 | *3.54*<br>73.3 | *4.05*<br>70.2 | *1.25*<br>17.9 |

**Supplementary Table 2.** Description of residue connectivity in EPA purified from Δ*epaR* cells[1] (EPA_epaR). This was determined by the analysis of $^{1}$H-$^{1}$H-NOESY and $^{1}$H-$^{13}$C-HMBC experiments from the anomeric shifts of each residue, recorded at 298 K.

| **Residue** | | **NOE** | **HMBC** |
|---|---|---|---|
| →3)-α-GlcNAc*p*-(1-*P* | **A** | None | None |
| →4)-6-*P*-α-MurNAc*p*-OH | **B** | None | None |
| α-Glc*p*-(1→ | **C** | H3, $\mathbf{R_H}$ | C3, $\mathbf{R_G}$ |
| α-Glc*p*-(1→ | **D** | H4, $\mathbf{R_D}$<br>H6, $\mathbf{R_D}$ | C4, $\mathbf{R_D}$ |
| →4)-6-*P*-β-MurNAc-OH | **E** | None | None |
| β-GlcNAc*p*-(1→ | **F** | None | None |
| β-GlcNAc*p*-(1→ | **G** | H2, $\mathbf{R_C}$ **or** $\mathbf{R_F}$ | C2, $\mathbf{R_C}$ |
| β-GlcNAc*p*-(1→ | **H** | H2, $\mathbf{R_A}$ | C2, $\mathbf{R_A}$ |
| β-GlcNAc*p*-(1→ | **H'** | H2, $\mathbf{R_C}$ **or** $\mathbf{R_F}$ | C2, $\mathbf{R_F}$ |
| β-GlcNAc*p*-(1→ | **I** | H4, **B or E** | C4, **B or E** |
| →2)-3-O-methyl-α-Rha*p*-(1→ | $\mathbf{R_A}$ | H3, $\mathbf{R_C}$ **or** $\mathbf{R_F}$ | C3, $\mathbf{R_C}$ |
| →2)-α-Rha*p*-(1→ | $\mathbf{R_B}$ | H3, $\mathbf{R_F}$ | C3, $\mathbf{R_F}$ |
| →2,3)-α-Rha*p*-(1→ | $\mathbf{R_C}$ | H2, $\mathbf{R_I}$ | C2, $\mathbf{R_I}$ **or** $\mathbf{R_E}$ |
| →2,4)-α-Rha*p*-(1→ | $\mathbf{R_D}$ | H2, $\mathbf{R_G}$ | C2, $\mathbf{R_G}$ |
| →2)-α-Rha*p*-(1→ | $\mathbf{R_E}$ | H3, $\mathbf{R_J}$ | C3, $\mathbf{R_J}$ |
| →2,3)-α-Rha*p*-(1→ | $\mathbf{R_F}$ | H2, $\mathbf{R_I}$ | C2, $\mathbf{R_I}$ **or** $\mathbf{R_E}$ |
| →2)-α-Rha*p*-(1→ | $\mathbf{R_G}$ | H3, $\mathbf{R_L}$ | C3, $\mathbf{R_L}$ |
| →2,3)-α-Rha*p*-(1→ | $\mathbf{R_H}$ | H2, $\mathbf{R_B}$ | C2, $\mathbf{R_B}$ **or** $\mathbf{R_F}$ |
| →2)-α-Rha*p*-(1→ | $\mathbf{R_I}$ | H2, $\mathbf{R_E}$ **or** $\mathbf{R_G}$ | C2, $\mathbf{R_E}$ **or** $\mathbf{R_G}$ |
| →2)-α-Rha*p*-(1→ | $\mathbf{R_J}$ | H2, $\mathbf{R_D}$ | C2, $\mathbf{R_D}$ |
| →3)-α-Rha*p*-(1→ | $\mathbf{R_K}$ | H2, $\mathbf{R_K}$ | C2, $\mathbf{R_K}$ **or** $\mathbf{R_B}$ |
| →3)-α-Rha*p*-(1→ | $\mathbf{R_L}$ | H3, **A** | C3, **A** |

1 Ho, K., Huo, W., Pas, S., Dao, R. & Palmer, K. L. Loss-of-Function Mutations in epaR Confer Resistance to φNPV1 Infection in *Enterococcus faecalis* OG1RF. *Antimicrob Agents Chemother* **62**, doi:10.1128/aac.00758-18 (2018).

**Supplementary Table 3.** All $^{1}H$ and $^{13}C$ NMR chemical shifts (ppm) at 298 K of EPA fragments (OS1 and OS2) produced following 18-hour treatment of EPA with hydrofluoric acid. Shifts were assigned through the analysis of $^{1}H$-$^{1}H$-COSY, $^{1}H$-$^{1}H$-TOCSY and $^{1}H$-$^{13}C$-HSQC experiments.

| **Residue** | | ***H1***<br>**C1** | ***H2***<br>**C2** | ***H3***<br>**C3** | ***H4***<br>**C4** | ***H5***<br>**C5** | ***H6***<br>**C6** | **H6'**<br>**C6** | **N-acetyl**<br>**$CH_3$** |
|---|---|---|---|---|---|---|---|---|---|
| **OS1** | | | | | | | | | |
| α-Glc*p*-(1→ | **J** | *4.92*<br>99.7 | *3.56*<br>72.7 | *3.73*<br>74.3 | *3.41*<br>70.9 | *3.67*<br>73.1 | *3.76*<br>61.8 | *3.86*<br>61.8 | |
| 1→)-Glycerol | **$Gro_1$** | *3.77/ 3.57*<br>69.8 | *3.96*<br>71.7 | *3.68/ 3.64*<br>63.75 | | | | | |
| Glycerol | **$Gro_2$** | *3.65/ 3.62*<br>63.7 | *ND*<br>*ND* | *3.57/ 3.55*<br>63.7 | | | | | |
| **OS2** | | | | | | | | | |
| α-GlcNAc*p*-(1→ | **K** | *5.23*<br>99.3 | *3.91*<br>55.1 | *3.79*<br>72.2 | *3.54*<br>70.9 | *4.02*<br>73.2 | *3.80*<br>61.5 | *ND*<br>61.5 | 2.03<br>23.4 |
| α-Glc*p*-(1→ | **L** | *4.97*<br>99.8 | *3.58*<br>72.6 | *3.71*<br>74.2 | *3.42*<br>70.9 | *3.69*<br>73.3 | *3.77*<br>61.8 | *3.88*<br>61.8 | |
| →3,6)-β-GalNAc*p*-(1→ | **M** | *4.76*<br>103.3 | *4.01*<br>52.7 | *3.92*<br>81.7 | *4.22*<br>69.5 | *3.87*<br>74.1 | *3.72*<br>68.7 | *3.96*<br>68.7 | 2.03<br>23.4 |
| →3)-β-GalNAc*p*-(1→ | **N** | *4.73*<br>103.3 | *4.00*<br>52.7 | *3.89*<br>81.8 | *4.19*<br>69.1 | *3.67*<br>76.2 | *3.7-3.9*<br>61.3- 62.5 | | 2.03<br>23.4 |
| →3,6)-β-Glc*p*-(1→ | **O** | *4.51*<br>106.0 | *3.40*<br>73.1 | *3.55*<br>83.4 | *3.70*<br>ND | *3.56*<br>75.9 | *3.83*<br>69.8 | *4.19*<br>69.8 | |
| →4)-β-GalNAc*p*-(1→ | **P** | *4.45*<br>103.5 | *3.88*<br>53.6 | *3.83*<br>72.5 | *4.13*<br>76.0 | *3.68*<br>75.6 | *3.88*<br>61.9 | *ND*<br>61.9 | 2.03<br>23.4 |
| β-Gal*p*-(1→ | **Q** | *4.43*<br>104.9 | *3.55*<br>72.1 | *3.66*<br>74.1 | *3.93*<br>70.0 | *3.69*<br>76.5 | *3.7-3.9*<br>61.3-62.5 | | |
| 1→)-Glycerol | **$Gro_3$** | *4.00/ 3.58*<br>71.9 | *3.84*<br>72.7 | *3.61/* 3.46<br>63.7 | | | | | |

**Supplementary Table 4.** Description of residue connectivity in EPA fragments (OS1 and OS2) produced following 18-hour treatment of EPA with hydrofluoric acid. This was determined by the analysis of $^{1}$H-$^{1}$H-ROESY and $^{1}$H-$^{13}$C-HMBC experiments from the anomeric shifts of each residue (or all shifts in the case of glycerol residues), recorded at 298 K.

| **Residue** | | **NOE** | **HMBC** |
|---|---|---|---|
| **OS1** | | | |
| α-Glc*p*-(1→ | **J** | H1 and H1', **Gro$_2$** | none |
| 1→)-Glycerol | **Gro$_1$** | none | none |
| Glycerol | **Gro$_2$** | none | none |
| **OS2** | | | |
| α-GlcNAc*p*-(1→ | **K** | H3, **O3** | C3, **O3** |
| α-Glc*p*-(1→ | **L** | H6 and H6', **M6** | none |
| →3,6)-β-GalNAc*p*-(1→ | **M** | H4, **P4** | C4, **P4** |
| →3)-β-GalNAc*p*-(1→ | **N** | H4, **P4** | C4, **P4** |
| →3,6)-β-Glc*p*-(1→ | **O** | H3, **M4 and N4** | C3, **M4 and N4** |
| →4)-β-GalNAc*p*-(1→ | **P** | H1, **Gro$_3$** | C1, **Gro$_3$** |
| β-Gal*p*-(1→ | **Q** | H6 and H6', **O6** | C6, **O6** |
| 1→)-Glycerol | **Gro$_3$** | none | none |

**Supplementary Table 5.** All $^{1}$H, $^{13}$C and $^{31}$P NMR chemical shifts (ppm) at 298 K of an EPA fragment (OS3) produced following 1-hour treatment of EPA with hydrofluoric acid. All shifts were determined through the analysis of $^{1}$H-$^{1}$H-COSY, $^{1}$H-$^{1}$H-TOCSY, $^{1}$H-$^{13}$C-HSQC, $^{1}$H-$^{13}$C-HSQC-TOCSY, $^{31}$P and $^{1}$H-$^{31}$P-HMQC-TOCSY experiments.

| **Residue** | | ***H1***<br>**C1** | ***H2***<br>**C2** | ***H3***<br>**C3** | ***H4***<br>**C4** | ***H5***<br>**C5** | ***H6***<br>**C6** | **H6'**<br>**C6** | **N-acetyl**<br>**$CH_3$** | **$PO_4$** |
|---|---|---|---|---|---|---|---|---|---|---|
| *P*-6-α-GlcNAc*p*-(1→ | **K*** | *5.27*<br>98.9 | *3.95*<br>55.0 | *3.80*<br>72.0 | *3.66*<br>70.4 | *4.14*<br>72.0 | *4.04*<br>65.2 | *4.19*<br>65.2 | *2.06*<br>23.6 | 0.98 |
| α-GlcNAc*p*-(1→ | **K** | *5.24*<br>99.1 | *3.92*<br>55.2 | *3.81*<br>71.9 | *3.55*<br>71.0 | *4.03*<br>73.0 | *3.79*<br>62.7 | *ND*<br>62.7 | *2.06*<br>23.6 | |
| α-Glc*p*-(1→ | **L** | *4.97*<br>99.7 | *3.56*<br>72.6 | *3.72*<br>74.3 | *3.42*<br>70.9 | *3.70*<br>73.1 | *3.78*<br>61.7 | *3.88*<br>61.7 | | |
| α-Glc*p*-(1→ | **R** | *4.95*<br>99.6 | *3.56*<br>72.6 | *3.76*<br>74.3 | *3.41*<br>70.9 | *3.70*<br>73.1 | *3.76*<br>61.7 | *3.87*<br>61.7 | | |
| →3,6)-β-GalNAc*p*-(1→ | **S*** | *4.72*<br>103.6 | *4.12*<br>52.1 | *3.89*<br>81.6 | *4.23*<br>69.3 | *3.92*<br>73.9 | *3.71*<br>69.6 | *3.96*<br>69.6 | *2.07*<br>23.6 | |
| →3)-β-GalNAc*p*-(1→ | **S** | *4.71*<br>103.7 | *4.12*<br>52.1 | *3.88*<br>81.7 | *4.19*<br>68.9 | *3.71*<br>76.1 | *3.84*<br>61.7 | *ND*<br>61.7 | *2.07*<br>23.6 | |
| →4)-*P*-3-β-GalNAc*p*-(1→ | **T** | *4.50*<br>103.1 | *3.97*<br>52.7 | *4.22*<br>76.2 | *4.28*<br>75.5 | *3.71*<br>75.4 | *3.81*<br>62.4 | *ND*<br>62.4 | *2.07*<br>23.6 | -0.51 |
| →3,6)-β-Glc*p*-(1→ | **O** | *4.53*<br>105.8 | *3.40*<br>73.0 | *3.57*<br>83.1 | *3.71*<br>71.1 | *3.57*<br>75.7 | *3.83*<br>69.4 | *4.19*<br>69.4 | | |
| β-Gal*p*-(1→ | **Q** | *4.43*<br>104.8 | *3.56*<br>72.0 | *3.65*<br>74.0 | *3.93*<br>69.9 | *3.69*<br>76.5 | *3.78*<br>62.4 | *3.81*<br>62.4 | | |
| *P*-3-β-Gal*p*-(1→ | **U** | *4.50*<br>104.4 | *3.70*<br>ND | *4.13*<br>79.1 | *4.13*<br>69.0 | *ND*<br>ND | *ND*<br>ND | *ND*<br>ND | | -0.21 |
| →1)-Glycerol | **Gro$_3$** | *3.59/ ND*<br>72.1 | *3.83*<br>71.8 | *3.60/ 3.55*<br>63.7 | | | | | | |
| Glycerol-3-*P* | **Gro$_4$** | *3.69/3.62*<br>63.4 | *3.65*<br>70.5 | *3.96/ 3.89*<br>67.8 | | | | | | 0.98 |
| →1)-Glycerol-3-*P* | **Gro$_5$** | *3.61/3.82*<br>69.5 | *4.09*<br>70.5 | *3.93/ ND*<br>67.6 | | | | | | -0.51 |

**Supplementary Table 6.** Description of residue connectivity in EPA fragment OS3 produced following 1-hour treatment of EPA with hydrofluoric acid. This was determined by the analysis of $^{1}$H-$^{1}$H-ROESY and $^{1}$H-$^{13}$C-HMBC experiments from the anomeric shifts of each residue (or all shifts in the case of glycerol residues), recorded at 298 K.

| **Residue** | | **NOE** | **HMBC** |
|---|---|---|---|
| *P*-6-α-GlcNAc*p*-(1→ | **K*** | H3, **O3** | C3, **O3** |
| α-GlcNAc*p*-(1→ | **K** | H3, **O3** | C3, **O3** |
| α-Glc*p*-(1→ | **L** | H6′, **S*6** | C6, **S*6** |
| α-Glc*p*-(1→ | **R** | H1 and H1′, **$Gro_5$** | C1, **$Gro_5$** |
| →3,6)-β-GalNAc*p*-(1→ | **S*** | H4, **T4** | H4, **T4** |
| →3)-β-GalNAc*p*-(1→ | **S** | H4, **T4** | H4, **T4** |
| →4)-*P*-3-β-GalNAc*p*-(1→ | **T** | H1 and H1′, **$Gro_3$** | C1, **$Gro_3$** |
| →3,6)-β-Glc*p*-(1→ | **O** | H3, **S3 and S*3** | C3, **S3 and S*3** |
| β-Gal*p*-(1→ | **Q** | H6 and H6′, **O6** | C6, **O6** |
| *P*-3-β-Gal*p*-(1→ | **U** | none | none |
| →1)-Glycerol | **$Gro_3$** | none | none |
| Glycerol-3-*P* | **$Gro_4$** | none | none |
| →1)-Glycerol-3-*P* | **$Gro_5$** | none | none |

**Supplementary Table 7.** All new or altered $^{1}H$, $^{13}C$ and $^{31}P$ NMR chemical shifts (ppm) at 298 K of EPA purified from Δ*11720* cells (EPA_11720; Smith et al., 2019) in reference against shifts observed in EPA_epaR (supplementary Table 1). All shifts were determined through the analysis of $^{1}H$-$^{1}H$-COSY, $^{1}H$-$^{1}H$-TOCSY, $^{1}H$-$^{13}C$-HSQC, $^{1}H$-$^{13}C$-HSQC-TOCSY, $^{31}P$ and $^{1}H$-$^{31}P$-HSQC-TOCSY experiments. No new phosphorus shifts were identified.

| **Residue** | | ***H1***<br>**C1** | ***H2***<br>**C2** | ***H3***<br>**C3** | ***H4***<br>**C4** | ***H5***<br>**C5** | ***H6***<br>**C6** | **H6'**<br>**C6** | **N-acetyl**<br>**$CH_3$** | **O-methyl**<br>**$CH_3$** |
|---|---|---|---|---|---|---|---|---|---|---|
| →2)-3-*O*-methyl-α-Rha*p*-(1→ | **$R_A$** | *5.27*<br>102.7 | *4.32*<br>75.9 | *3.41*<br>80.0 | *3.42*<br>72.2 | *3.65*<br>70.5 | 1.31<br>18.1 | | | *3.43*<br>57.3 |
| 3→)-β-GlcNAc*p*-(1→ | **H** | *4.61*<br>103.9 | *3.74*<br>56.1 | *3.64*<br>80.9 | 3.73<br>73.2 | *3.48*<br>75.4 | *3.62*<br>61.1 | *ND*<br>61.1 | *2.08*<br>23.6 | |
| α-Glc*p*-(1→ | **V** | *4.94*<br>99.4 | *3.54*<br>72.6 | *3.78*<br>73.9 | *3.39*<br>70.8 | *3.66*<br>73.1 | *3.86*<br>61.7 | *ND*<br>61.7 | | |
| →6)-β-GalNAc*p*-(1→ | **W** | *4.53*<br>103.2 | *3.95*<br>53.4 | *3.77*<br>71.8 | *3.98*<br>68.8 | *3.97*<br>74.5 | *3.97*<br>67.7 | *3.66*<br>67.7 | *2.08*<br>23.6 | |

**Supplementary Table 8.** Description of connectivity of residues described in Supplementary Table 7 corresponding to EPA_11720. This was determined by the analysis of $^{1}H$-$^{1}H$-NOESY and $^{1}H$-$^{13}C$-HMBC experiments from the anomeric shifts of each residue, recorded at 298 K.

| **Residue** | | **NOE** | **HMBC** |
|---|---|---|---|
| →2)-3-*O*-methyl-α-Rha*p*-(1→ | **R$_A$** | H3, **R$_c$ or R$_F$** | C3, **R$_c$** |
| 3→)-β-GlcNAc*p*-(1→ | **H** | H2, **R$_A$** | C2, **R$_A$** |
| α-Glc*p*-(1→ | **V** | H6 and H6’, **W** | C6, **W** |
| →6)-β-GalNAc*p*-(1→ | **W** | H3, **H** | C3, **H** |

**Supplementary Table 9.** All new or altered $^{1}$H, $^{13}$C and $^{31}$P NMR chemical shifts (ppm) at 298 K of EPA purified from Δ*11715* cells (EPA_11715; Smith et al., 2019) in reference against shifts observed in EPA_11720 (supplementary Table 7). All shifts were determined through the analysis of $^{1}$H-$^{1}$H-COSY, $^{1}$H-$^{1}$H-TOCSY, $^{1}$H-$^{13}$C-HSQC, $^{1}$H-$^{13}$C-HSQC-TOCSY, $^{31}$P and $^{1}$H-$^{31}$P-HSQC-TOCSY experiments.

| **Residue** | | ***H1***<br>**C1** | ***H2***<br>**C2** | ***H3***<br>**C3** | ***H4***<br>**C4** | ***H5***<br>**C5** | ***H6***<br>**C6** | **H6'**<br>**C6** | **N-acetyl**<br>**CH$_3$** | **PO$_4$** |
|---|---|---|---|---|---|---|---|---|---|---|
| *P*-6-α-GlcNAc*p*-(1→ | **K*** | *5.24*<br>98.9 | *3.96*<br>55.9 | *3.77*<br>72.0 | *3.63*<br>70.4 | *4.14*<br>71.9 | *4.15*<br>65.1 | *4.03*<br>65.1 | *2.07*<br>23.5 | 1.05 |
| α-GlcNAc*p*-(1→ | **K** | *5.22*<br>99.1 | *3.93*<br>55.0 | *3.77*<br>72.0 | *ND*<br>*ND* | *ND*<br>*ND* | *ND*<br>*ND* | *5.22*<br>99.1 | *2.07*<br>23.5 | |
| →3,6)-β-GalNAc*p*-(1→ | **W*** | *4.59*<br>102.9 | *4.06*<br>52.3 | *3.90*<br>81.0 | *4.23*<br>68.9 | *ND*<br>*ND* | *3.98*<br>67.7 | *3.68*<br>67.7 | *2.06*<br>23.5 | |
| →3)-β-GalNAc*p*-(1→ | **W** | *4.58*<br>102.9 | *4.04*<br>52.3 | *3.88*<br>81.5 | *4.26*<br>68.8 | *ND*<br>*ND* | *3.63-3.90*<br>61.0-61.9 | | *2.06*<br>23.5 | |
| β-Glc*p*-(1→ | **O** | *4.51*<br>105.6 | *3.29*<br>74.0 | *3.46*<br>76.7 | *3.41*<br>70.6 | *ND*<br>*ND* | *3.87*<br>61.7 | *3.73*<br>61.7 | | |
| →3)-β-Glc*p*-(1→ | **O*** | *4.53*<br>105.6 | *3.38*<br>72.8 | *3.56*<br>82.9 | *3.60*<br>71.1 | *ND*<br>*ND* | *3.87*<br>61.7 | *3.73*<br>61.7 | | |
| Glycerol-3-*P* | **Gro$_4$** | *3.68 3.62*<br>63.3 | *3.90*<br>71.8 | *3.95/3.88*<br>67. 6 | | | | | | 1.05 |

**Supplementary Table 10.** Description of connectivity of residues described in Supplementary Table 9 corresponding to EPA_11715. This was determined by the analysis of $^{1}H$-$^{1}H$-NOESY and $^{1}H$-$^{13}C$-HMBC experiments from the anomeric shifts of each residue (or all shifts in the case of glycerol residues), recorded at 298 K.

| **Residue** | | **NOE** | **HMBC** |
|---|---|---|---|
| *P*-6-α-GlcNAc*p*-(1→ | **K*** | H3, **O*3** | C3, **O*3** |
| α-GlcNAc*p*-(1→ | **K** | H3, **O*3** | C3, **O*3** |
| →3,6)-β-GalNAc*p*-(1→ | **W*** | H3, **H3** | C3, **H3** |
| →3)-β-GalNAc*p*-(1→ | **W** | H3, **H3** | C3, **H3** |
| β-Glc*p*-(1→ | **O** | H3, **W3 and W*3** | C3, **W3 and W*3** |
| →3)-β-Glc*p*-(1→ | **O*** | H3, **W3 and W*3** | C3, **W3 and W*3** |
| Glycerol-3-*P* | **$Gro_4$** | none | none |

**Supplementary Table 11.** All new or altered $^{1}H$, $^{13}C$ and $^{31}P$ NMR chemical shifts (ppm) at 298 K of EPA purified from Δ*11714* cells (EPA_11714; Smith et al., 2019) in reference against shifts observed in EPA_11715 (supplementary Table 9). All shifts were determined through the analysis of $^{1}H$-$^{1}H$-COSY, $^{1}H$-$^{1}H$-TOCSY, $^{1}H$-$^{13}C$-HSQC, $^{1}H$-$^{13}C$-HSQC-TOCSY, $^{31}P$ and $^{1}H$-$^{31}P$-HSQC-TOCSY experiments.

| **Residue** | | ***H1*** **C1** | ***H2*** **C2** | ***H3*** **C3** | ***H4*** **C4** | ***H5*** **C5** | ***H6*** **C6** | **H6’ C6** | **$PO_4$** |
|---|---|---|---|---|---|---|---|---|---|
| →3,6)-β-Glc*p*-(1→ | **O*** | *4.53* 105.7 | *3.37* 73.0 | *3.56* 82.6 | *3.69* 71.1 | *3.56* 75.5 | *4.19* 69.5 | *3.85* 69.5 | |
| →6)-β-Glc*p*-(1→ | **O** | *4.52* 105.7 | *3.31* 74.0 | *3.46* 76.7 | *3.69* 71.1 | *3.56* 75.5 | *4.21* 69.9 | *3.84* 69.9 | |
| *P*-3-β-Gal*p*-(1→ | **U** | *4.50* 104.2 | *3.68* 71.1 | *4.09* 79.0 | *4.13* 69.0 | *3.71* 76.1 | *ND* ND | *ND* ND | -0.10 |
| Glycerol-3-*P* | **$Gro_7$** | *3.66/ 3.58* 63.4 | *4.06* 70.8 | *3.98/3.91* 67.4 | | | | | -0.10 |

**Supplementary Table 12.** Description of connectivity of residues described in Supplementary Table 11 corresponding to EPA_11714. This was determined by the analysis of $^{1}H$-$^{1}H$-NOESY and $^{1}H$-$^{13}C$-HMBC experiments from the anomeric shifts of each residue (or all shifts in the case of glycerol residues), recorded at 298 K.

| **Residue** | | **NOE** | **HMBC** |
|---|---|---|---|
| →3,6)-β-Glc*p*-(1→ | **O*** | H3, **W and W*** | C3, **W and W*** |
| →6)-β-Glc*p*-(1→ | **O** | H3, **W and W*** | C3, **W and W*** |
| *P*-3-β-Gal*p*-(1→ | **U** | H6 and H6*, **O and O*** | C6, **O and O*** |
| Glycerol-3-*P* | **Gro$_4$** | none | none |

**Supplementary Table 13.** All new or altered $^{1}H$, $^{13}C$ and $^{31}P$ NMR chemical shifts (ppm) at 298 K of EPA purified from Δ*11706* cells (EPA_11706) in reference against shifts observed in EPA_11714 (supplementary Table 11). All shifts were determined through the analysis of $^{1}H$-$^{1}H$-COSY, $^{1}H$-$^{1}H$-TOCSY, $^{1}H$-$^{13}C$-HSQC, $^{1}H$-$^{13}C$-HSQC-TOCSY, $^{31}P$ and $^{1}H$-$^{31}P$-HSQC-TOCSY experiments.

| **Residue** | | ***H1*** **C1** | ***H2*** **C2** | ***H3*** **C3** | ***H4*** **C4** | ***H5*** **C5** | ***H6*** **C6** | **H6’ C6** | **N-acetyl $CH_3$** | **$PO_4$** |
|---|---|---|---|---|---|---|---|---|---|---|
| α-Glc*p*-(1→ | **J** | *4.94*<br>*99.6* | *3.56*<br>72.7 | *3.76*<br>74.2 | *3.41*<br>70.9 | *3.70*<br>73.11 | *3.86*<br>61.8 | *3.76*<br>61.8 | | |
| →3,6)-β-GalNAc*p*-(1→ | **S*** | *4.71*<br>*103.7* | *4.13*<br>52.1 | *3.88*<br>81.3 | *4.20*<br>69.3 | *3.92*<br>73.9 | *3.95*<br>68.5 | *3.70*<br>68.5 | *2.07*<br>23.8 | |
| →3)-β-GalNAc*p*-(1→ | **S** | *4.691*<br>*103.7* | *4.13*<br>52.1 | *3.88*<br>81.3 | *4.18*<br>68.9 | *3.71*<br>76.1 | *3.84*<br>61.7 | *ND*<br>61.7 | *2.07*<br>23.8 | |
| →4)-*P*-3-β-GalNAcp-(1→ | **T** | *4.51*<br>103.4 | *3.97*<br>52.9 | *4.21*<br>76.1 | *4.27*<br>75.8 | *3.71*<br>75.3 | *3.81*<br>62.4 | *ND*<br>62.4 | *2.07*<br>23.8 | -0.45 |
| *P*-3-β-Gal*p*-(1→ | **U** | *4.48*<br>104.4 | *3.68*<br>71.1 | *4.01*<br>78.9 | *4.13*<br>69.0 | *3.71*<br>76.1 | *ND*<br>ND | *ND*<br>ND | | -0.14 |
| →1)-Glycerol-3-*P* | **$Gro_5$** | *3.61/3.82*<br>69.4 | *4.08*<br>70.4 | *3.95/3.93*<br>67.5 | | | | | | -0.45 |
| →1)-Glycerol-3-*P* | **$Gro_6$** | *3.97/3.63*<br>72.2 | *4.00*<br>70.7 | *3.94/3.84*<br>67.5 | | | | | | -0.14 |

**Supplementary Table 14.** Description of connectivity of residues described in Supplementary Table 13 corresponding to EPA_11706. This was determined by the analysis of $^{1}$H-$^{1}$H-NOESY and $^{1}$H-$^{13}$C-HMBC experiments from the anomeric shifts of each residue (or all shifts in the case of glycerol residues), recorded at 298 K.

| **Residue** | | **NOE** | **HMBC** |
|---|---|---|---|
| α-Glc*p*-(1→ | **J** | H1 and H1', **Gro$_5$** | C1, **Gro$_5$** |
| →3,6)-β-GalNAc*p*-(1→ | **S*** | H4, **T** | C4, **T** |
| →3)-β-GalNAc*p*-(1→ | **S** | H4, **T** | C4, **T** |
| →4)-*P*-3-β-GalNAcp-(1→ | **T** | H1 and H1', **Gro$_6$** | C1, **Gro$_6$** |
| *P*-3-β-Gal*p*-(1→ | **U** | H6 and H6', **O and O*** | C6, **O and O*** |
| →1)-Glycerol-3-*P* | **Gro$_5$** | none | none |
| →1)-Glycerol-3-*P* | **Gro$_6$** | none | none |

**Supplementary Table 15.** Bioinformatic analysis of *epaA*-*epaQ* in *E. faecalis* OG1RF

| Protein | Structural homologue[a] | | | | |
|---|---|---|---|---|---|
| | **Protein** | **Organism** | **Activity** | **TM-score** | **Reference** |
| **EpaA** | WecA | *Mycobacterium tuberculosis* H37Rv | Decaprenyl-phosphate *N*-acetylglucosaminephosphotransferase | 0.79 | (1) |
| **EpaB** | WbbL | *Mycobacterium tuberculosis* H37Rv | *N*-acetylglucosaminyl-diphospho-decaprenol L-rhamnosyltransferase | 0.86 | (2) |
| **EpaC** | RhlC | *Pseudomonas aeruginosa* PA01 | α-1,2-rhamnosyltransferase | 0.81 | (3) |
| **EpaD** | WapR | *Pseudomonas aeruginosa* PA01 | α-1,3-rhamnosyltransferase | 0.67 | (4) |
| **EpaE** | Cps2L | *Streptococcus pneumoniae* R6 | Glucose-1-phosphate thymidylyltransferase | 0.99 | (5) |
| **EpaF** | RfbC | *Escherichia coli* K12 | dTDP-4-dehydrorhamnose 3,5-epimerase | 0.92 | (6) |
| **EpaG** | RmlB | *Mycobacterium tuberculosis* H37Rv | dTDP-glucose 4,6-dehydratase | 0.99 | (7, 8) |
| **EpaH** | RmlD | *Mycobacterium tuberculosis* H37Rv | dTDP-4-dehydrorhamnose reductase | 0.89 | (9) |
| **EpaI** | GacI | *Streptococcus pyogenes* | Decaprenyl β -*N*-acetylglucosaminephosphotransferase | 0.93 | (10) |
| **EpaJ** | GacJ | *Streptococcus pyogenes* | Forms a complex with GacI which increases GacI activity | 0.96 | (10) |
| **EpaK** | None | | | | |
| **EpaL:EpaM**[b] | Wzm:Wzt | *Aquifex aeolicus* VF5 | ABC transporter complex | 0.54 | (11) |
| **EpaN** N-terminal | Predicted (*DW089_08855*) | *Acidaminococcus sp.* AM05-11 | Class I SAM-dependent methyltransferase | 0.70 | N/A |
| **EpaN** C-terminal | WsaE[c] | *Geobacillus stearothermophilus* NRS 2004/3a | α-1,2 and α-1,3- rhamnosyltransferase | 0.82 | (12) |
| **EpaO** | WsaE[c] | *Geobacillus stearothermophilus* NRS 2004/3a | α-1,2 and α-1,3- rhamnosyltransferase | 0.91 | (12) |
| **EpaP** | Predicted (*arnT1*) | *Pseudomonas fluorescens* Pf-5 | Undecaprenyl phosphate-α-4-amino-4 deoxy-L-arabinose arabinosyl transferase 1 | 0.63 | N/A |
| **EpaQ** | Predicted (*KPHS_51260*) | *Klebsiella pneumoniae subsp. Pneumoniae* HS11286 | Predicted *O*-antigen polymerase | 0.62 | N/A |

[a] Structural homologues were identified either directly in the literature (GacI and GacJ) or by comparison of AlphaFold 2 structures (13), proteins with TM-scores of above 0.5 were deemed to be structural homologues (14).

[b] The formation of the the EpaL:EpaM complex was predicted using AlphaFold-Multimer (15)

[c] Following the removal of the N-terminal 2-*O*-methlytransferase domain of WsaE (12)

1 Jin Y, Xin Y, Zhang W, Ma Y. *Mycobacterium tuberculosis* Rv1302 and *Mycobacterium smegmatis* MSMEG_4947 have WecA function and MSMEG_4947 is required for the growth of *M. smegmatis*. FEMS Microbiol Lett. 2010;310(1):54-61.

2 Mills JA, Motichka K, Jucker M, Wu HP, Uhlik BC, Stern RJ, et al. Inactivation of the mycobacterial rhamnosyltransferase, which is needed for the formation of the arabinogalactan-peptidoglycan linker, leads to irreversible loss of viability. J Biol Chem. 2004;279(42):43540-6.

3 Rahim R, Ochsner UA, Olvera C, Graninger M, Messner P, Lam JS, et al. Cloning and functional characterization of the *Pseudomonas aeruginosa* rhlC gene that encodes rhamnosyltransferase 2, an enzyme responsible for di-rhamnolipid biosynthesis. Mol Microbiol. 2001;40(3):708-18.

4 Poon KK, Westman EL, Vinogradov E, Jin S, Lam JS. Functional characterization of MigA and WapR: putative rhamnosyltransferases involved in outer core oligosaccharide biosynthesis of *Pseudomonas aeruginosa*. J Bacteriol. 2008;190(6):1857-65.

5 Jakeman DL, Young JL, Huestis MP, Peltier P, Daniellou R, Nugier-Chauvin C, et al. Engineering ribonucleoside triphosphate specificity in a thymidylyltransferase. Biochemistry. 2008;47(33):8719-25.

6 Macpherson DF, Manning PA, Morona R. Characterization of the dTDP-rhamnose biosynthetic genes encoded in the rfb locus of *Shigella flexneri*. Mol Microbiol. 1994;11(2):281-92.

7 Li W, Xin Y, McNeil MR, Ma Y. rmlB and rmlC genes are essential for growth of mycobacteria. Biochem Biophys Res Commun. 2006;342(1):170-8.

8 Tsukioka Y, Yamashita Y, Oho T, Nakano Y, Koga T. Biological function of the dTDP-rhamnose synthesis pathway in *Streptococcus mutans*. J Bacteriol. 1997;179(4):1126-34.

9 Tsukioka Y, Yamashita Y, Nakano Y, Oho T, Koga T. Identification of a fourth gene involved in dTDP-rhamnose synthesis in *Streptococcus mutans*. J Bacteriol. 1997;179(13):4411-4.

10 Rush JS, Edgar RJ, Deng P, Chen J, Zhu H, van Sorge NM, et al. The molecular mechanism of N-acetylglucosamine side-chain attachment to the Lancefield group A carbohydrate in *Streptococcus pyogenes*. J Biol Chem. 2017;292(47):19441-57.

11 Spellmon N, Muszynski A, Gorniak I, Vlach J, Hahn D, Azadi P, et al. Molecular basis for polysaccharide recognition and modulated ATP hydrolysis by the O antigen ABC transporter. Nat Commun. 2022;13(1):5226.

12 Steiner K, Novotny R, Werz DB, Zarschler K, Seeberger PH, Hofinger A, et al. Molecular basis of S-layer glycoprotein glycan biosynthesis in *Geobacillus stearothermophilus*. J Biol Chem. 2008;283(30):21120-33.

(13) Jumper J, Evans R, Pritzel A, Green T, Figurnov M, Ronneberger O, et al. Highly accurate protein structure prediction with AlphaFold. Nature. 2021;596(7873):583-9.

14 Zhang Y, Skolnick J. TM-align: a protein structure alignment algorithm based on the TM-score. Nucleic Acids Research. 2005;33(7):2302-9.

15 Evans R, O'Neill M, Pritzel A, Antropova N, Senior A, Green T, et al. Protein complex prediction with AlphaFold-Multimer. bioRxiv. 2022:2021.10.04.463034.

**Supplementary Table 16.** Bioinformatic analysis of *E. faecalis epaR* and downstream genes

| Protein | Structural homologue[a] | | | | |
|---|---|---|---|---|---|
| | Protein | Organism | Activity | TM-score | Reference |
| **EpaR** | WecP | *Aeromonas hydrophila* AH-3 | UDP-N-acetylgalactosamine-undecaprenyl-phosphate N-acetylgalactosaminephosphotransferase | 0.67 | (1) |
| **11720** | WfgD | *Escherichia coli* O152 | β-1,3-glucosyltransferases | 0.91 | (2) |
| **11719** | | | N/A[b] | | |
| **11718** | | | N/A | | |
| **11717** | Orf2 | *Streptococcus pneumoniae* R6 | Transposase | 0.61 | (3) |
| **11716** | | | N/A | | |
| **11715** | CgtB | *Campylobacter jejuni* OH4384 | β-1,3-galactosyltransferase activity | 0.81 | (4) |
| **11714** | CgtB | *Campylobacter jejuni* OH4384 | β-1,3-galactosyltransferase activity | 0.83 | (4) |
| **11713** | TarF | *Staphylococcus aureus* PS 47 | Glycerol-phosphate transferase | 0.84 | (5) |
| **11712** | TarD | *Staphylococcus aureus* PS 47 | Glycerol-3-phosphate cytidylyltransferase | 0.95 | (6) |
| **11711** | TarF | *Staphylococcus aureus* PS 47 | Glycerol-phosphate transferase | 0.84 | (5) |
| **11710** | WaaL | *Shigella dysenteriae* | Lipid A-core:surface polymer ligase | 0.69 | (7) |
| **11709** | Cps2J | *Streptococcus pneumoniae* R6 | Wzx transporter (flippase) | 0.94 | (8) |
| **11708** | | | N/A | | |
| **11707** | GalE1 | *Mycobacterium tuberculosis* CDC 1551 | UDP-glucose 4-epimerase | 0.93 | (9) |
| **11706** | CsbB | *Bacillus subtilis* 168 | Decaprenyl α-N-acetylglucosaminephosphotransferase | 0.91 | (10) |

[a] Structural homologues were identified by comparison of AlphaFold 2 structures (11); proteins with TM-scores of above 0.5 were deemed to be structural homologues (12).

[b] N/A, no activity (truncated protein or no homolog detected)


1 Merino S, Jimenez N, Molero R, Bouamama L, Regue M, Tomas JM. A UDP-HexNAc:polyprenol-P GalNAc-1-P transferase (WecP) representing a new subgroup of the enzyme family. J Bacteriol. 2011;193(8):1943-52.

2 Brockhausen I, Hu B, Liu B, Lau K, Szarek WA, Wang L, et al. Characterization of two beta-1,3-glucosyltransferases from *Escherichia coli* serotypes O56 and O152. J Bacteriol. 2008;190(14):4922-32.

3 Orihuela CJ, Radin JN, Sublett JE, Gao G, Kaushal D, Tuomanen EI. Microarray analysis of pneumococcal gene expression during invasive disease. Infect Immun. 2004;72(10):5582-96.

4 Gilbert M, Brisson JR, Karwaski MF, Michniewicz J, Cunningham AM, Wu Y, et al. Biosynthesis of ganglioside mimics in Campylobacter jejuni OH4384. Identification of the glycosyltransferase genes, enzymatic synthesis of model compounds, and characterization of nanomole amounts by 600-mhz (1)h and (13)c NMR analysis. J Biol Chem. 2000;275(6):3896-906.

5 Brown S, Zhang YH, Walker S. A revised pathway proposed for *Staphylococcus aureus* wall teichoic acid biosynthesis based on in vitro reconstitution of the intracellular steps. Chem Biol. 2008;15(1):12-21.

6 Badurina DS, Zolli-Juran M, Brown ED. CTP:glycerol 3-phosphate cytidylyltransferase (TarD) from *Staphylococcus aureus* catalyzes the cytidylyl transfer via an ordered Bi-Bi reaction mechanism with micromolar K(m) values. Biochim Biophys Acta. 2003;1646(1-2):196-206.

7 Ruan X, Loyola DE, Marolda CL, Perez-Donoso JM, Valvano MA. The WaaL O-antigen lipopolysaccharide ligase has features in common with metal ion-independent inverting glycosyltransferases. Glycobiology. 2012;22(2):288-99.

8 Xayarath B, Yother J. Mutations blocking side chain assembly, polymerization, or transport of a Wzy-dependent *Streptococcus pneumoniae* capsule are lethal in the absence of suppressor mutations and can affect polymer transfer to the cell wall. J Bacteriol. 2007;189(9):3369-81.

9 Skovierova H, Larrouy-Maumus G, Pham H, Belanova M, Barilone N, Dasgupta A, et al. Biosynthetic origin of the galactosamine substituent of Arabinogalactan in *Mycobacterium tuberculosis*. J Biol Chem. 2010;285(53):41348-55.

10 Rismondo J, Percy MG, Grundling A. Discovery of genes required for lipoteichoic acid glycosylation predicts two distinct mechanisms for wall teichoic acid glycosylation. J Biol Chem. 2018;293(9):3293-306.

11 Jumper J, Evans R, Pritzel A, Green T, Figurnov M, Ronneberger O, et al. Highly accurate protein structure prediction with AlphaFold. Nature. 2021;596(7873):583-9.

12 Zhang Y, Skolnick J. TM-align: a protein structure alignment algorithm based on the TM-score. Nucleic Acids Research. 2005;33(7):2302-9.

**Supplementary table S17: Bacterial strains, plasmids, and primers.**

| Strains/plasmids/primers | Relevant properties/sequence [a] | Source |
|---|---|---|
| **Strains** | | |
| *Enterococcus faecalis* | | |
| OG1RF | Plasmid-free, virulent strain isolated from a human oral cavity | 1 |
| OG1RF Δ*11706* | OG1RF with an in-frame deletion of *OG1RF_11706* | This work |
| OG1RF Δ*11706* GFP | OG1RF Δ*11706* expressing GFP from pMV158-GFP; Tet5 | This work |
| OG1RF Δ*11706* (pTet-11706) GFP | OG1RF Δ*11706* mutant complemented and expressing GFP; Tet5, Erm30 | This work |
| OG1RF Δ*11714* | OG1RF with an in-frame deletion of *OG1RF_11714* | 2 |
| OG1RF Δ*11714* GFP | OG1RF Δ*11714* expressing GFP from pMV158-GFP; Tet5 | This work |
| OG1RF Δ*11714* (pTet-11706) GFP | OG1RF Δ*11714* mutant complemented and expressing GFP; Tet5, Erm30 | This work |
| OG1RF Δ*11715* | OG1RF with an in-frame deletion of *OG1RF_11715* | 2 |
| OG1RF Δ*11715* GFP | OG1RF Δ*11715* expressing GFP from pMV158-GFP; Tet5 | This work |
| OG1RF Δ*11715* (pTet-11706) GFP | OG1RF Δ*11715* mutant complemented and expressing GFP; Tet5, Erm30 | This work |
| OG1RF Δ*11720* | OG1RF with an in-frame deletion of *OG1RF_11720* | 2 |
| OG1RF Δ*11720* GFP | OG1RF Δ*11720* expressing GFP from pMV158-GFP; Tet5 | This work |
| OG1RF Δ*11720* (pTet-11706) GFP | OG1RF Δ*11706* mutant complemented and expressing GFP; Tet5, Erm30 | This work |
| OG1RF Δ*epaR* | OG1RF with an in-frame deletion of *OG1RF_11721* | 3 |
| OG1RF Δ*epaR::epaR* | OG1RF Δ*epaR* complemented mutant (knock in) | 3 |
| *Escherichia coli* | | |
| TG1 (*repA*$^{+}$) | TG1 derivative encoding RepA for pGhost9 propagation at 37°C | 4 |
| **Plasmids** | | |
| pGhost9 | Temperature-sensitive plasmid for gene replacement; Erm30 | 5 |
| pGhost-11706 | pGhost9 derivative used to build the in-frame deletion of *OG1RF_11706*; Erm30 | This work |
| pMV158-GFP | pMV158 derivative expressing the *gfp* gene under control of a constitutive promoter; Tet5. | 6 |
| pTetH | pAT18 derivative encoding TetR for tetracycline-inducible expression in *E. faecalis;* Erm30 | [7] |
| pTet-11706 | pTetH derivative to complement the *OG1RF_11706* deletion; Erm30 | This work |
| pTet-11714 | pTetH derivative to complement the *OG1RF_11714* deletion; Erm30 | 2 |
| pTet-11715 | pTetH derivative to complement the *OG1RF_11715* deletion; Erm30 | 2 |
| pTet-11720 | pTetH derivative to complement the *OG1RF_11720* deletion; Erm30 | 2 |
| **Oligonucleotides** | | |
| SM_0171 (pGhost9_up) | GTCACGACGTTGTAAAACGACGG | |
| SM_0172 (pGhost9_down) | CTAGCGGACTCTAGAGGATCCCA | |
| SM_0216 (11706_H11) | CCCCTCGAGTCCATTAACGCCTTATGCAGTGG | |
| SM_0217 (11706_H12) | CGGCCAACATACTCCCCATTATTTGCTTCATTATACGCGGGTATTG | |
| SM_0218 (11706_H21) | TAATGAAGCAAATAATGGGGAGTATGTTGGCCGTGTAT | |
| SM_0219 (11706_H22) | GTGGCGGCCGCGGCAATTTCTGTCTTAGGATCAGC | |
| SM_0256 (11706_H110) | TTGTACCATTGGAGTGAGGTGGG | |
| SM_0257 (11706_H220) | CGAGCTGGGGACATCAAAGATTCC | |
| SM_0100 (pTetH_Fw) | GCTTGATCGTAGCGTTAACAGATCTACTC | |
| SM_0101 (pTetH_Rev) | CAAATTGTGGATGTGACCATGCGG | |

| | |
|---|---|
| SM_0348 (11706_Fw) | AGTGAGCTCAAGGAGGAGACTGACCATGGGGAAAAAGAAAATTTTAATTTCAATACCCGCGTATAATGAAG |
| SM_0349 (11706_Rev) | GTGGGATCCATCAATCTCTTTTAACTCTGATTCTTTAGTCTCATTTTTC |

[a] Tet5, Tetracycline 5µgml$^{-1}$; Erm30, Erythromycin 30µg ml$^{-1}$
